# Probing PAC1 receptor activation across species with an engineered sensor

**DOI:** 10.1101/2024.02.06.579048

**Authors:** Reto B. Cola, Salome N. Niethammer, Preethi Rajamannar, Andrea Gresch, Musadiq A. Bhat, Kevin Assoumou, Elyse Williams, Patrick Hauck, Nina Hartrampf, Dietmar Benke, Miriam Stoeber, Gil Levkowitz, Sarah Melzer, Tommaso Patriarchi

## Abstract

Class-B1 G protein-coupled receptors (GPCRs) are an important family of clinically relevant drug targets that remain difficult to investigate via high-throughput screening and in animal models. Here, we engineered PAClight1_P78A_, a novel genetically-encoded sensor based on a class-B1 GPCR (the human PAC1 receptor, hmPAC1R) endowed with high dynamic range (ΔF/F_0_ = 1100%), excellent ligand selectivity and rapid activation kinetics (τ_ON_ = 1.15 sec). To showcase the utility of this tool for in vitro applications, we thoroughly characterized and compared its expression, brightness and performance between PAClight1_P78A_ transfected and stably-expressing cells. Demonstrating its use in animal models, we show robust expression and fluorescence responses upon exogenous ligand application ex vivo and in vivo in mice, as well as in living zebrafish larvae. Thus, the new GPCR-based sensor can be used for a wide range of applications across the life sciences empowering both basic research and drug development efforts.

## Introduction

Class-B1 G protein-coupled receptors (GPCRs) represent an important sub-group of peptide-sensing GPCRs, that are the focus of intense and rapidly-expanding drug development efforts^1^, driven by extremely successful examples of peptide agonists used in the clinical treatment of metabolic human diseases, such as type-2 diabetes and obesity^2^. One such peptide-GPCR system that has shown growing potential for targetability in the treatment of human disorders is the Pituitary Adenylate Cyclase Activating Peptide (ADCYAP1 or PACAP) and its receptors. PACAP is an endogenous 38-amino acid peptide that is amongst the most phylogenetically-conserved peptides^3^. Its shorter C-terminally-truncated form (i.e. PACAP_1-27_) has 68% homology with the vasoactive-intestinal-peptide (VIP)^4^. In fact, VIP and PACAP share a subfamily of class-B1 G protein-coupled receptors (GPCRs), of which the two receptors VPAC1 and VPAC2 can both be equipotently activated by VIP and PACAP^4^. The third receptor in this subfamily, i.e. the PAC1 receptor (PAC1R, also known as ADCYAP1R1), however, has a reported affinity for PACAP that is 100-1000-fold higher than its affinity to VIP^4^. The tissue distribution of PACAP and its receptors is widespread and they can be found throughout the central and peripheral nervous system, the immune system, in endocrine glands as well as in other organ systems and in many cancerous tissues^4–8^. A large body of evidence has linked the PACAP/PAC1R system to protective functions in the nervous and immune systems, as well as to stress- and anxiety-related behaviors (in particular post-traumatic stress disorder (PTSD)), migraine, nociception, thermoregulation, sleep/wake cycles, and reproductive functions^4,7,9^, making it a peptide signaling system of high clinical relevance.

Recently, genetically-encoded GPCR-based sensors have been developed that enable the direct optical detection of GPCR activation by agonist ligands with high sensitivity and spatiotemporal resolution^10–18^. While these tools hold great potential for drug development, pharmacology and neuroscience applications, only a few of them are built from class-B1 peptide-sensing GPCRs^17,19^. Thus, the development of new biosensors based on peptide-sensing GPCRs would greatly benefit the community and serve as a powerful resource for drug screening and life sciences.

In this work we engineered PAClight1_P78A_, an ultrasensitive indicator based on the human PAC1 receptor (hmPAC1R). To establish this as a tool for peptide drug screening, we thoroughly characterized its dynamic range, as well as its optical, kinetic, signaling and pharmacological properties in vitro. We further generated stable cell lines and compared them to transfected cells for ligand-induced fluorescence responses using flow cytometry. Additionally, we tested the potential of PAClight1_P78A_ as a tool to study ligand binding and diffusion in animal models. To this end, we examined the sensitivity and specificity of PAClight1_P78A_ in acute mouse brain slices in response to application of PACAP_1-38_. Moreover, we verified ligand detection in vivo using fiber photometry recording and intracerebral microinfusions in behaving mice.

Finally, we demonstrate that the sensor expresses well and produces a large fluorescent response to PACAP_1-38_ following microinjection in the brain of living zebrafish larvae, opening new opportunities for the development and testing of drugs targeting this receptor in the central nervous system.

Overall, our new sensor expands the class of genetically-encoded optical tools that can be used for GPCR-targeted HTS assays, reducing the demands on time and costly reagents, and provides new opportunities for functionally testing drugs that target the PAC1R pathway directly in living animals.

## Results

### Development of an ultrasensitive PAC1R-based sensor

To develop a highly-sensitive indicator of hmPAC1R activation, we followed a protein engineering approach that we and others recently established^10,13,16^. We used the human PAC1Rnull splice isoform as a protein scaffold (hmPAC1R) and constructed an initial sensor prototype in which we replaced the entire third intracellular loop (ICL3, residues Q336-G342) with a module containing circularly-permuted green fluorescent protein (cpGFP) from dLight1.3b^10^ (**Extended Data Fig. 1a**). This initial prototype construct, named PAClight0.1, was expressed on the surface of HEK293T cells, but showed clear intracellular retention and a very weak average fluorescent response to bath application of PACAP_1-38_ (ΔF/F_0_ = 43.4%, **Extended Data Fig. 1b**). Using this sensor as template, we conducted a small-scale screen to identify the optimal insertion site by reintroducing amino acids from the original ICL3 of hmPAC1R on both sides of the cpGFP-module. This led us to the identification of a second mutant, in which Q336 was reintroduced before the cpGFP-module, that showed improved average fluorescent response to the ligand (ΔF/F_0_ = 343%, **Extended Data Fig. 1c**). To improve membrane expression of the PAClight mutants, we next investigated the effect of point mutations on their C-terminus, via site-directed mutagenesis. Mutation into alanine of residue T468, a reported post-translational modification (PTM) site (www.phosphosite.org) on the C-terminus of the receptor^20^, improved the overall surface expression of the sensor in HEK293T cells and further increased the observed average fluorescent response (ΔF/F_0_ = 516%, **Extended Data Fig. 1f**). To further improve the fluorescent response of PAClight, we screened a subsequent library of sensor variants with mutations targeted to the second intracellular loop (ICL2). A similar approach was previously shown to boost the fluorescent response of other GPCR-based sensors, for example dLight1.3b or OxLight1^10,16^. Our functional screen of a library of ICL2 mutants led to the identification of one variant comprising three point mutations (F259K/F260K/P261K) that displayed a largely improved fluorescent response to PACAP_1-38_ (ΔF/F_0_ = 942%, **Extended Data Fig. 1d**). Upon introduction of the C-term T468A mutation on the ICL2 F259K/F260K/P261K background, we obtained an improved version of the sensor, named PAClight1, with excellent surface expression and fluorescent response (ΔF/F_0_ = 1037%, **Extended Data Fig. 1g**). Given the high degree of sequence and structural similarity, as well as the broad activation of endogenous VPAC receptors by vasoactive intestinal peptide (VIP) and PACAP^21^, we next asked whether the hmPAC1R-based sensor would also respond to VIP. Indeed, application of a high concentration of VIP onto sensor-expressing cells caused a large fluorescent response, corresponding to more than half of the response to PACAP_1-38_ (ΔF/F_0_ = 654%, **Extended Data Fig. 2**c and 2d). In an effort to eliminate the response to VIP and obtain a PACAP-specific sensor, we screened a small library of sensors containing single-point mutations into alanine that were inspired by simulations on the binding free energy of PACAP and VIP on the PAC1R^22^. Furthermore, at this stage we included the naturally occurring splice mutant PAC1R ‘short’ (reported to have lower affinity towards VIP^8^) into the extracellular ligand-binding domain (ECD) of the sensor, as well as two structure-guided point-mutants. Through this screening, we identified a single mutation (P78A) that abolished the sensor response to VIP while leaving unaltered the response to PACAP_1-38_ (**Extended Data Fig. 2c and 2d**). The final sensor construct, named PAClight1_P78A_ (**Figure 1a,b**), includes all the above-mentioned mutations and combines a very large dynamic range with excellent PACAP selectivity and good membrane expression, and was thus selected for further *in vitro* characterization.

**Figure 1.**
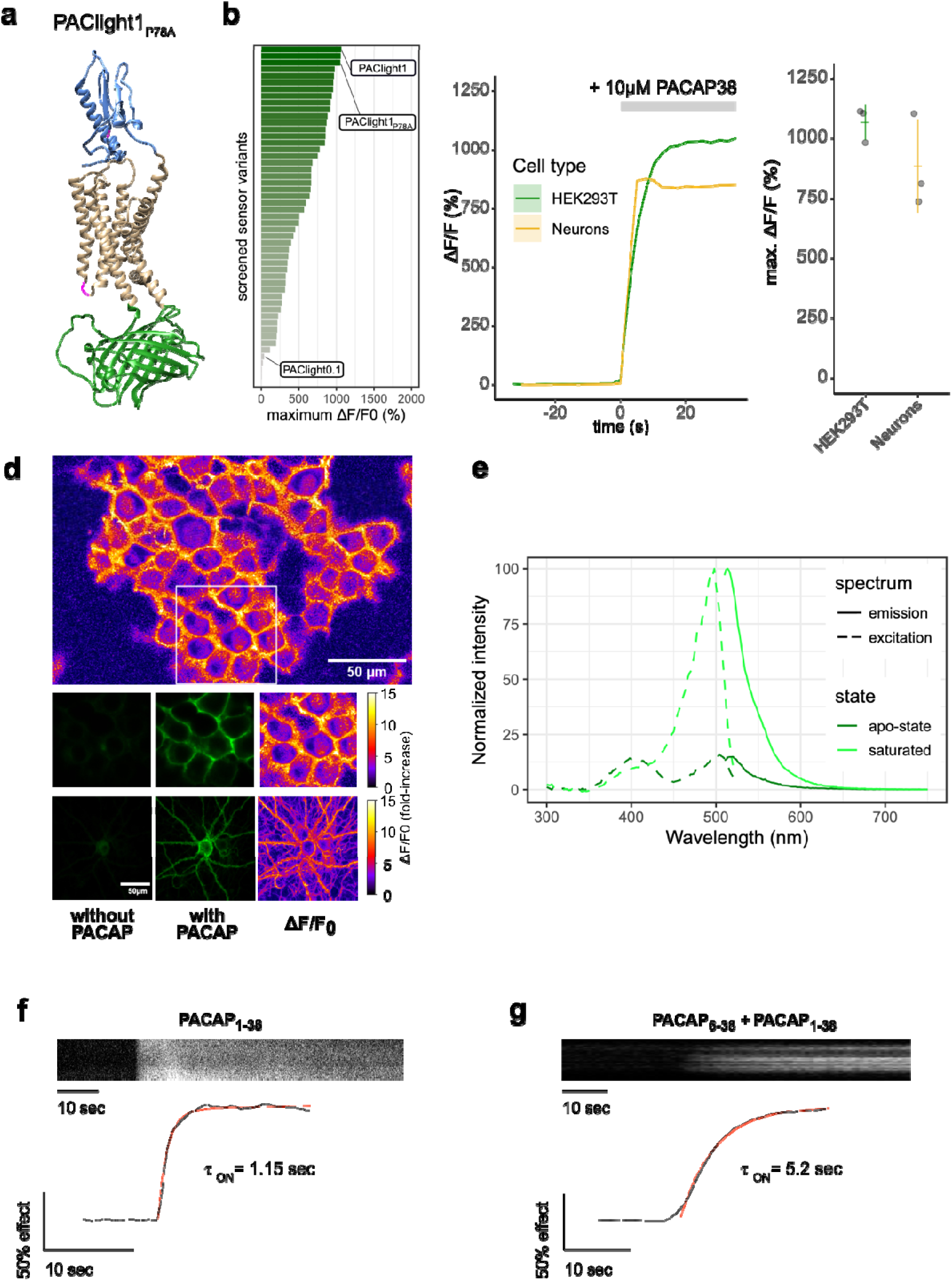
Design and in vitro optical properties of PAClight1_P78A_. **a)** Protein structure of PAClight1_P78A_ as predicted by AlphaFold2^50^. The transmembrane and intracellular domain of the PAC1Rnull backbone is depicted in beige. The extracellular domain, which is crucial for ligand specificity and affinity, is colored in blue. The cpGFP module replacing the 3^rd^ intracellular loop is colored in green. Point-mutations inserted in the 2^nd^ intracellular loop as well as on the extracellular domain are depicted in magenta. **b)** Bar chart depicting the maximum dynamic range obtained for all tested mutants with acceptable expression on the plasma membrane. Bars are ordered by dynamic range and color coded by the average of the maximum dynamic range recorded. The bars representing the prototype variant PAClight_V0.1, PAClight1_P78A_ and PAClight1 are indicated. **c)** Line plots depicting the maximum activation (mean +/−SEM) of PAClight1_P78A_ expressed in HEK293T cells and rat primary neurons. Upon bath application of 10 μM PACAP_1-38,_ PAClight1_P78A_ reaches a mean ΔF/F_0_ of 1066% in HEK293T cells (n=3, 5 ROIs) and a mean ΔF/F_0_ of 883% in rat primary neurons (*n*=3, 5ROIs). **c’)** Scatter plot representation of the maximum ΔF/F_0_ of the individual replicates shown in **c**. Confocal image acquisition was performed at a frame rate of 1 frame/2.53s (∼0.4Hz). **d)** Representative examples of the expression of PAClight1_P78A_ in HEK293T cells (low magnification overview and middle row of images) and rat primary neurons (bottom row) with pixel-wise quantification and depiction of the dynamic range in ΔF/F_0_ upon bath application of 10μM PACAP_1-38_. Top: Overview micrograph of HEK293T cells color coded in pixel-wise ΔF/F_0_. The white square represents the ROI represented below in the middle row of images. Middle: Selected ROI before (left) and after (middle and right) peptide application. Bottom: Rat primary neuron expressing PAClight1_P78A_ before (left) and after (middle and right) peptide application. The color bars represent the look-up table used for visualization of the pixel-wise ΔF/F_0_. **e)** One-photon excitation (dashed lines) and emission (solid lines) spectra of PAClight1_P78A_ in the absence (apo-state, dark green) and the presence (saturated state, light green) of 10μM PACAP_1-38_. The excitation maximum of the saturated state is at 498nm. The isosbestic point is at 420nm. Emission maximum in of the saturated state is at 514nm. *n*=4 (2 replicates each measured on 2 independent days). **f**) Activation kinetics of PAClight1_P78A_ upon application of PACAP_1-38_ (10 μM) measured via timelapse imaging. A representative kymograph of sensor fluorescence on the surface of a HEK293 cell is shown on top. The normalized fluorescence response trace is shown at bottom along with the calculated one-phase association curve fit and activation time constant. The trace shown is the average of 3 independent experiments. **g**) Same as **f)** but in the presence of PACAP_6-38_ (10 μM) in the bath.

### In vitro characterization of PAClight1_P78A_

The newly developed PAClight1_P78A_ sensor displays an average fluorescent response of 1066% ΔF/F_0_ in HEK293T cells (n=3, 5 ROIs each) to bath application of 10μM PACAP_1-38_ (**Figure 1b-d**). We verified that sensor expression and function is not drastically affected by the cell type in which it is expressed by testing it in primary cultured neurons. Neurons were virally transduced using an adeno-associated virus (AAV) for expressing the PAClight1_P78A_ sensor under control of a human synapsin-1 promoter. 2-3 weeks after transduction we verified excellent expression of the probe on the plasma membrane of the neurons with no appreciable intracellular retention. Under these conditions, the sensor showed a fluorescence response of 883% ΔF/F_0_ upon bath application of the PACAP_1-38_ ligand (**Figure 1c-d**).

We next set out to determine the sensor’s excitation and emission spectra in vitro in HEK29T cells (**Figure 1e**). The excitation maxima in the absence and presence of 10μM PACAP_1-38_ were identified at 504nm and 498nm respectively. The isosbestic point at which the excitation is independent of the absence or presence of PACAP_1-38_ is located at 420nm. The maxima for the emission spectra in the absence of PACAP_1-38_ was identified at 520nm and in the presence of PACAP_1-38_ at 514nm.

Our previous work on the development of another class-B1 sensor based on the GLP1 receptor, led us to discover that the kinetics of the sensor’s response can be used to infer the occupancy of the ECD by an antagonist peptide^17^. Given that all class-B1 GPCRs share a similar ECD high-affinity ligand binding mechanism, we performed similar experiments to determine whether PAClight1_P78A_ could also be used in a similar manner. To do so, we monitored the sensor’s response during application of PACAP_1-38_ alone or in the presence of PACAP_6-38_, an antagonist peptide, in the buffer surrounding the cells. We then determined the activation time constant of the fluorescent response in both conditions. The PAClight1_P78A_ response was strikingly slower (approximately 4-fold) in the presence of the extracellular antagonist, and was in the range of 1 second in its absence **(Figure 1f,g)**. Thus, the kinetics of PACligth1_P78A_ response could be used to investigate or screen for factors that influence the ECD-PACAP interaction and potentially the speed of signal transduction through conformational activation of the receptor.

To investigate the stability of the fluorescent response to bath application of PACAP_1-38_, a long-term imaging experiment was performed at room temperature with 1 frame (1024×1024 pixels, 2x line-averaging) acquired every minute over the time course of 150 min (**Figure 2a**). After bath application of 200nM PACAP_1-38_ (slow diffusion of the peptide with 10x dilution from 2μM to 200nM) there was no decrease in signal, nor any internalization observed. After >100 frames of acquisition. we applied a saturating bolus of the peptide PAC1R-antagonist Max.d.4. Within 45-50 frames after application of the PAC1R antagonist, the signal intensity steadily decreased and started to plateau slightly above baseline levels. This indicates, that Max.d.4 can outcompete PACAP_1-38_ at PACligth1_P78A_, albeit not to a full extent under the specified experimental conditions.

**Figure 2.**
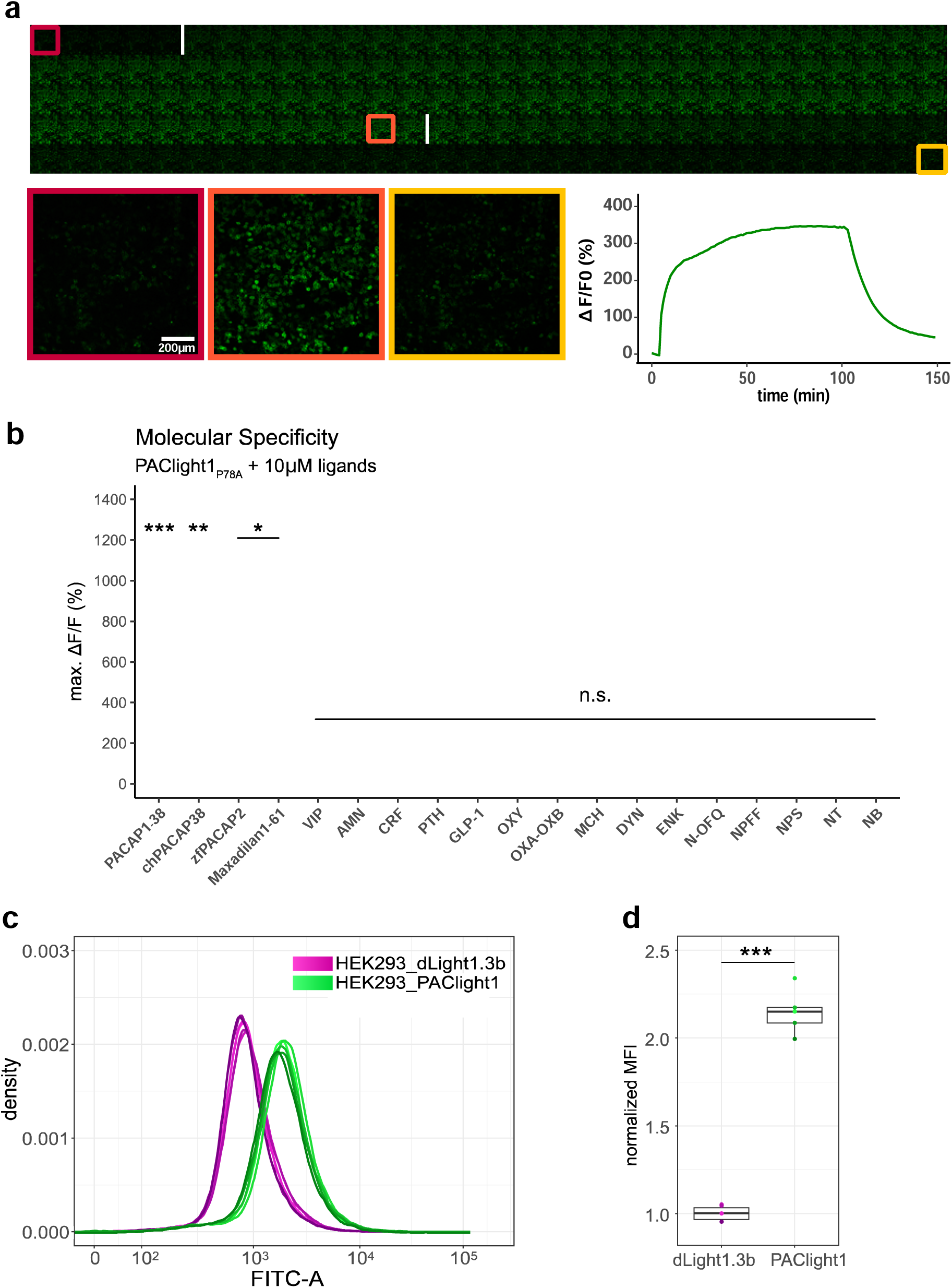
Pharmacological characterization of PAClight1_P78A_r. **a)** PAClight1_P78A_ fluorescent response to 200 nM PACAP**_1-38_** bath application over extended periods at room temperature. No internalization of the sensor expressed on the plasma membrane is observed throughout the full time course of >90 minutes. The fluorescent response of PAClight1_P78A_ to PACAP**_1-38_** can be reversed with competitive binding of the peptidergic PAC1R antagonist Max.d.4 (a Maxadilan derivative). Please note that the seemingly slow activation rate of PAClight1_P78A_ in this experiment is due to the experimental setup (see methods section) and slow diffusion of PACAP_1-38_ throughout the well. Top: Downscaled gallery view of the acquired time series (total of 150 frames; 1 frame/minute) from top-left to bottom-right. Application time points of PACAP**_1-38_** (after frame Nr. 5) and Max.d.4 (after frame Nr. 103) are depicted as white vertical lines between the frames. The red, orange, and yellow rectangles indicate the frames used for representative higher magnification inserts shown below (bottom left). Bottom right: Line plot depicting the time course of the PAClight1_P78A_ response across 5 rectangular ROIs distributed across the whole field of view. **b)** PAClight1_P78A_ is highly specific to PACAP and does not respond to VIP. PAClight1_P78A_ can be activated by PACAP homologues found in the chicken (gallus gallus, chPACAP38) as well as in the zebrafish (danio rerio, zfPACAP2). PAClight1_P78A_ is further partially activated by the PAC1R specific ligand Maxadilan**_1-61_**, which is expressed endogenously in the sand fly (Lutzomyia longipalpis) salivary gland. None of the other tested class A and B1 GPCR ligands was found to activate PAClight1_P78A_. Abbreviations: AMN amnesiac, CRF corticotropin-releasing factor, PTH parathyroid hormone, GLP-1 glucagon-like peptide, OXY oxytocin, OXA-OXB orexin-A and -B, MCH melanin-concentrating hormone, DYN dynorphin, ENK enkephalin, N-OFQ nociception, NPFF neuropeptide FF, NPS neuropeptide S, NT neurotensin, NB neuromedin B. Single data points represent one replicate average obtained from 5 ROIs per replicate. The extent of the colored vertical bar represents 1 standard deviation. The y-axis location of the colored horizontal bar indicates the average across all replicates. Number of replicates per ligand: n=5 for chPACAP38 and GLP-1; n=4 for PACAP**_1-38_**, zfPACAP2, and Maxadilan**_1-61_**; n=3 for VIP, AMN, CRF, PTH, OXY, OXA-OXB, MCH, DYN, ENK, N-OFQ, NPS, NT, and NB; n=2 for NPFF. Asterisks represent statistical significance of Hochberg-corrected *p* values of multiple one-sample *t* tests. **c)** PAClight1 has significantly brighter baseline fluorescence than the dopamine sensor dLight1.3b. The peak of the dLight1.3b FITC-A density curve coincides with the start of the uphill slope of the PAClight1 FITC-A density curve. Note the bi-exponential scaling of the x-axis. n=6 for dLight1.3b, n=5 for PAClight1. **d)** Quantification of the average of the median fluorescence intensity (MFI) across all replicates and normalization to the group MFI of dLight1.3b show a 2.15-fold increased basal brightness of PAClight1 over dLight1.3b (*t*_(4.74)_= - 19.254, *p* < 0.0001, 95% CI [−1.30, −0.99], two-sided two-sample Welch’s *t* test).

Next, we screened a range of different peptides at saturating concentrations (10μM) for potential activation of the PAClight1_P78A_ sensor (**Figure 2b**). Mammalian PACAP_1-38_ displayed the strongest potency in PAClight1_P78A_ activation (1052% ΔF/F_0_, *t*_(3)_= 40.31, 95% CI [9.69, 11.35], *p*<0.001). Because the amino acid sequence of PACAP is rather well conserved throughout phylogeny, we also tested chicken PACAP_1-38_ (chPACAP38), as well as the zebrafish PACAP2_1-27_ (zfPACAP2) for potency on the PAClight1_P78A_ sensor. Both of these homologues of the mammalian PACAP_1-38_ activated the PAClight1_P78A_ sensor with strong but slightly reduced potency (chPACAP38: 896% ΔF/F_0_, *t*_(4)_= 11.17, 95% CI [6.73, 11.19], *p*= 0.0066; zfPACAP2: 898% ΔF/F_0_, *t*_(3)_= 11.49, 95% CI [6.46, 11.41], *p*= 0.024). Furthermore, we also screened the sand fly salivary gland derived peptide Maxadilan, which was previously identified to be a specific ligand of the PAC1R but not to the VPAC1 and VPAC2 receptors^23^. Consistent with the reported activity of Maxadilan on the hmPAC1Rnull receptor, we detected activation of PAClight1_P78A_ by Maxadilan, however, with lower potency than mammalian PACAP_1-38_ (416% ΔF/F_0_, *t*_(3)_= 9.47, 95% CI [2.76, 5.56], *p*= 0.04). The reduced potency of Maxadilan for PAClight1_P78A_ might be a consequence of the ligand specificity determining point-mutation (P78A) introduced into the extracellular domain of the PAClight1_P78A_ sensor. The drosophila gene *amn* (amnesiac) was previously shown to be homologous to the mammalian gene encoding PACAP (*Adcyap1*) and *amn* drosophila mutants show memory impairments similar to rodent PACAP/PAC1R mutants^24^. We therefore also tested whether this insect peptide encoded by *amn* would activate PAClight1_P78A_, but no significant activation above baseline was observed (26% ΔF/F_0_, *t*_(2)_= 3.2, 95% CI [−0.09, 0.61], *p*= 0.69). As shown above (**Extended Data Fig. 2c and 2d**) PAClight1_P78A_ was specifically optimized for VIP non-responsive properties. Therefore, the response of PAClight1_P78A_ to VIP in this specificity screen was also not significantly above baseline (57% ΔF/F_0_, *t*_(2)_= 11.09, 95% CI [0.35, 0.79], *p*= 0.12). Other peptides tested in this screen included other class B1 GPCR ligands (corticotropin-releasing factor (CRF), parathyroid hormone (PTH), and glucagon-like peptide 1 (GLP-1)), as well as some peptidergic class A GPCR ligands (oxytocin (OXY), orexin-A and -B (OXA-OXB), melanin-concentrating hormone (MCH), dynorphin (DYN), enkephalin (ENK), nociception (N-OFQ), neuropeptide FF (NPFF), neuropeptide S (NPS), neurotensin (NT), and neuromedin B (NB)). None of these neuropeptides activated the PAClight1_P78A_ sensor above baseline level (**Figure 2b**). Taken together, these results highlight the broad potential applicability of the PAClight1_P78A_ sensor for use in model systems across the phylogenetic tree, as well as its high selectivity for PACAP ligands over VIP and other peptide GPCR ligands.

In order not to induce artificial PACAP signaling and potentially interfere with downstream readouts when using PAClight1_P78A_, it is important to verify that the sensor does not recruit G-proteins and/or β-arrestin. To monitor the capacity of PACLight1_P78A_ to engage these intracellular signaling partners, we performed split NanoLuc complementation assays as in our previous work^16,18^, using either PAClight1_P78A_-SmBiT or, as a positive control, PAC1R-SmBiT fusion constructs, together with LgBiT-miniGs (**Extended Data Fig. 3a**), LgBiT-miniGsq (**Extended Data Fig. 3b**), or LgBiT-β-arrestin2 (**Extended Data Fig. 3c**). As expected, we observed significant miniGs, miniGsq, and β-arrestin2 recruitment to the wild-type PAC1R upon activation with 1 μM of PACAP_1-38_. Yet, we did not detect recruitment of either miniGs, miniGsq, or β-arrestin2 in PAClight1_P78A_-expressing cells upon stimulation with PACAP_1-38_. These results indicate that expression of PAClight1_P78A_ does not artificially induce PACAP-mediated intracellular signaling and is not likely to interfere with endogenous signaling pathways.

### Development of non-responsive PAClight1 control sensors

When employing GPCR-sensors in intact living tissue (e.g., when used in animal models) it is often desirable to make use of an appropriate control sensor, in which ligand-binding is abolished by virtue of one or more point mutations in the GPCR binding pocket. To engineer such control sensors for our PAClight1_P78A_ and PAClight1 sensors, we targeted key residues in the hmPAC1R that interact with residue D3 in PACAP. The carboxylic group of D3 was shown to be crucial for binding affinity and biological activity of PACAP to all three receptor types (i.e. PAC1, VPAC1, and VPAC2)^25,26^. Furthermore, the cryo-EM structure of the hmPAC1R in interaction with PACAP identified residues Y161 and R199 of the hmPAC1R to interact with residue D3 of PACAP^21^. Residue Y161 forms a hydrogen bond with D3, while R199 forms an electrostatic interaction with D3 (**Extended Data Fig. 4a and 4b**). To abolish these interactions, we first mutated R199 into alanine (R199A) on the backbone of PAClight1. This single point mutant showed drastically reduced average fluorescent response (PAClight1: 1075% ΔF/F_0_, PAClight1_R199A: 41.9% ΔF/F_0_, **Extended Data Fig. 4c**) to 10μM PACAP_1-38_. To further abolish the remaining response, we additionally mutated Y161 into alanine (Y161A) on the previous backbone (i.e. PAClight1_R199A_Y161A) and named the construct “PAClight1-ctrl”. This completely reduced the fluorescent response to PACAP_1-38_ to baseline levels (PAClight1-ctrl: 2.87% ΔF/F_0_). Next, we cloned these two point mutations into the backbone of PAClight1_P78A_ and observed an equally abolished fluorescent response to PACAP_1-38_ (PAClight1_P78A_: 1034% ΔF/F_0_, PAClight1_P78A_-ctrl: 10.44% ΔF/F_0_, *t*_(4.03)_ = 37.89, *p* < 0.0001, 95% CI [1098.63, 949.02], two-sided two-sample Welch’s *t* test, **Extended Data Fig. 4d**) in transfected HEK293T cells, as well as in virally-transduced rat primary neuron cultures (**Extended Data Fig. 4f**). No drastic differences in basal brightness were observed between PAClight1_P78A_ and PAClight1_P78A_-ctrl constructs (**Extended Data Fig. 4e**). In summary, we have developed double point-mutant sensor constructs for both PAClight1_P78A_ and PAClight1 with fully abolished responses to bath application of saturating concentrations of PACAP_1-38_.

### Comparison between transient and stable expression of the sensor

During the development and validation of PAClight1_P78A_, we noticed higher basal brightness levels of PAClight1_P78A_ compared to other GPCR-based fluorescent sensors. To obtain a quantitative comparison, we decided to produce a stable T-Rex HEK293 cell line for inducible expression of PAClight1 (mutant missing P78A ECD mutation). We then used flow cytometry to record multiple replicates of 100,000 cells, and compared their fluorescence intensity to that of a similarly generated cell line expressing the indicator dLight1.3b, which we previously described^27^ (**Figure 2c-d**). Visualization of the fluorescent readouts clearly showed a shift of the density curves towards higher fluorescence for HEK293_PAClight1 cells compared to the HEK293_dLight1.3b. Statistical comparison of the distributions of the two cell lines shows 2.15-fold higher basal brightness of PAClight1 over dLight1.3b (*t*_(4.74)_= −19.254, *p* < 0.0001, 95% CI [−1.30, −0.99], two-sided two-sample Welch’s *t* test, **Figure 2d**).

Among the most important potential advantages of using stable cell lines expressing GPCR sensors for pharmacological assays are the homogeneity of expression, reproducibility, and ease of use. To verify that this is indeed the case, we analyzed PACLight1_P78A_ expressing stable cells and compared them to transfected cells using flow cytometry. While a large proportion of cells from the transfected condition expressed the sensor at very low levels or not at all (i.e., were left shifted in the density plot), we also observed a significant proportion of very bright and strongly expressing cells from within the same condition. In fact, there is higher abundance of very bright and strongly expressing cells in the transfected condition than in the induced stable cell line condition (**Figure 3a**). The combination of the higher abundance of very low expression levels and very high expression levels within the transfected condition leads to a significantly increased variability of observed expression levels compared to the induced stable cell line condition. The median of the standard deviation in the FITC-A channel across the titration series was 62.6% smaller in the stable PAClight1_P78A_ relative to the transfected PAClight1_P78A_ condition (*p*= 0.0142, **Figure 3b**). The variability of cells transfected with PAClight1 was very comparable and was only 8.6% lower than in the PAClight1_P78A_ transfected condition (*p*= 0.5966, **Figure 3b**). The difference of 8.6% (even though statistically unsignificant) might partially be explained by slightly different performance of PAClight1_P78A_ vs. PAClight1 in dynamic range. *P* values represent the result of Dunnett’s test for correction of multiple comparison that was performed after the statistically significant results of an omnibus ANOVA (*F*_(2,30)_= 4.321, *p*= 0.0224).

**Figure 3.**
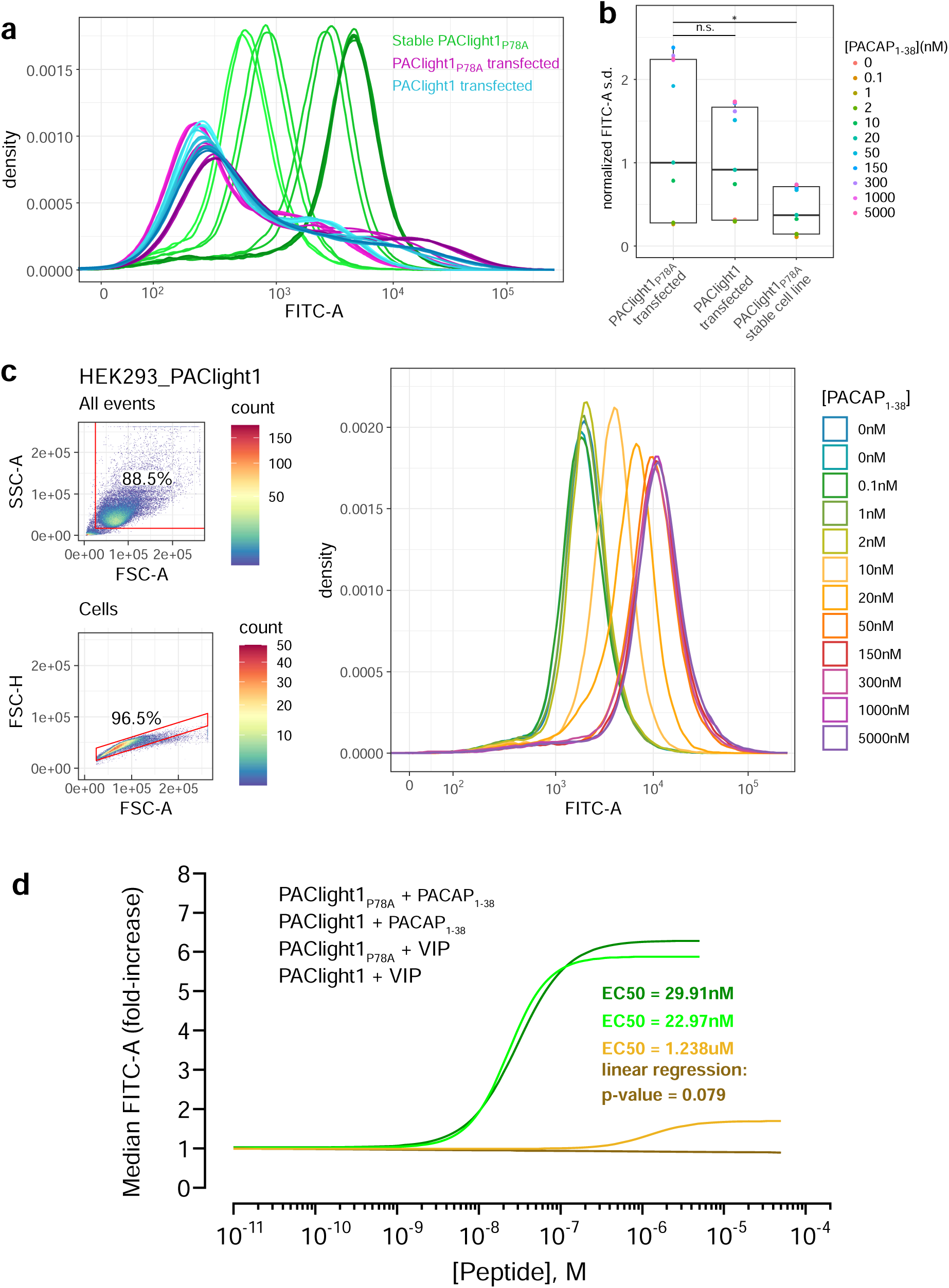
Comparison between transient and stable PAClight1_P78A_ expression. **a)** A direct flow cytometric comparison between transfected HEK293T cells and stable and inducible HEK293_PAClight1_P78A_ cells highlights the improved homogeneity of the distribution of expression levels in the newly generated stable cell line. Note the increased number of cells on both the low and high extremes in the transfected populations over the stable cell line population. Data from a full PACAP_1-38_ titration series for each condition is presented. The overall variability in expression levels in the PAClight1_P78A_- and PAClight1-transfected condition is very similar. **b)** The standard deviation of each density curve (normalized to the median standard deviation of the transfected PAClight1_P78A_ condition) is plotted by condition and concentration. The median standard deviation of the stable HEK293_PAClight1_P78A_ condition is reduced to only 37.4% of the transfected PAClight1_P78A_ condition (Dunnett’s test *p*= 0.0142). The median standard deviation varies little between transfected PAClight1_P78A_ and PAClight1 cells (8.6% lower in PAClight1, Dunnett’s test *p*= 0.5966). **c)** A representative example of a PACAP_1-38_ titration series on stable HEK293_PAClight1 cells. Top left: Gating strategy used to gate cells by the area of the side scatter (SSC-A) vs. the area of the forward scatter (FSC-A). Bottom left: Gating strategy used to gate singlets (within the cells gate) by the height of the forward scatter (FSC-H) vs. the FSC-A. Right: Density curves of the FITC-A channel obtained from the singlet gate. **d)** Dose-response curves obtained from VIP and PACAP_1-38_ titrations on stable HEK293_PAClight1_P78A_ and stable HEK293_PAClight1 cells. PAClight1_P78A_ and PAClight1 both have high affinities for PACAP_1-38_ (PAClight1_P78A_ EC50 = 29.91nM, PAClight1 EC50 = 22.97nM, n=3). While PAClight1 still shows a response to higher concentrations of VIP (EC50 = 1.24μM), PAClight1_P78A_’s response to VIP is completely abolished up to concentrations of 50μM VIP (*F*_(1, 33)_ = 3.28, *p*= 0.079, adj. R2= 0.06). Data in **a-c)** is derived from 100K original events recorded for each concentration and construct. Data in **d)** is derived from 100K recorded events across n=3 of each titration series.

With the stable cells and their improved homogeneity of expression levels, we then determined the affinities of PAClight1_P78A_ and PAClight1 towards PACAP_1-38_ and VIP. For each peptide concentration of a titration series 100,000 events were acquired, gated for cell clusters and for singlets. The FITC-A median fluorescence intensity (MFI) of the events in the singlet gate were used for dose-response fitting. A representative PACAP_1-38_ titration series replicate on stable HEK293_PAClight1 cells is shown in **Figure 3c**. The titration dataset reveals a slightly lower affinity of the PAClight1_P78A_ sensor (EC50= 29.91nM) than the PAClight1 sensor (EC50 = 22.97nM) towards PACAP_1-38_ (**Figure 3d**). The affinity of the PAClight1 sensor (non-specific mutant) towards VIP (EC50 = 1.238μM) is much lower than its affinity towards PACAP_1-38_ (**Figure 3d**). This 100-1000x difference is well in accordance with previous reports on the differential affinities of VIP and PACAP_1-38_ on the PAC1R^28,29^. We were not able to detect any response of PAClight1_P78A_ to the addition of VIP up to a concentration of 50μM (linear regression *p*= 0.079, **Figure 3d**), corroborating the specificity of the PAClight1_P78A_ sensor. Taken together, these data show that the PAClight1_P78A_ and PAClight1 sensors display very high affinity towards PACAP_1-38_, with a slightly lower affinity of PAClight1 compared to PAClight1_P78A_ as the result of the point mutation on the extracellular domain of PAClight1_P78A_.

### Characterization of PAClight1_P78A_ and PAClight1_P78A_-ctrl in model organism systems

Since PAClight1_P78A_ showed excellent expression and response properties in neuronal cell cultures, we next tested whether the sensor can be established as a tool to test ligand binding in intact neuronal circuits of mammalian model systems. Publicly available RNAseq databases (*in situ* hybridization atlas from the Allen Institute: alleninstitute.org) and previous work^30^ show strong expression of PACAP receptor (*Adcyap1r1*) in the cerebral cortex and hippocampus of mice and humans, suggesting these structures as interesting candidates for drug targeting. We tested the expression, sensitivity, and specificity of PAClight1_P78A_ in acute mouse brain slices that provide a physiological environment to examine the actions of pharmacological agents within specific brain areas^31^.

AAVs encoding the PAClight1_P78A_ or PAClight1_P78A_-ctrl sensor under control of a human synapsin-1 promoter were stereotactically injected into the neocortex and hippocampus of adult mice (**Figure 4a**). 4 weeks later, virus expression in the injection site was validated by histology (**Figure 4b**). PAClight1_P78A_ and PAClight1_P78A_-ctrl were efficiently expressed in neuronal cell bodies, axons and dendrites (**Figure 4c, Extended Data Figure 5**).

**Figure 4.**
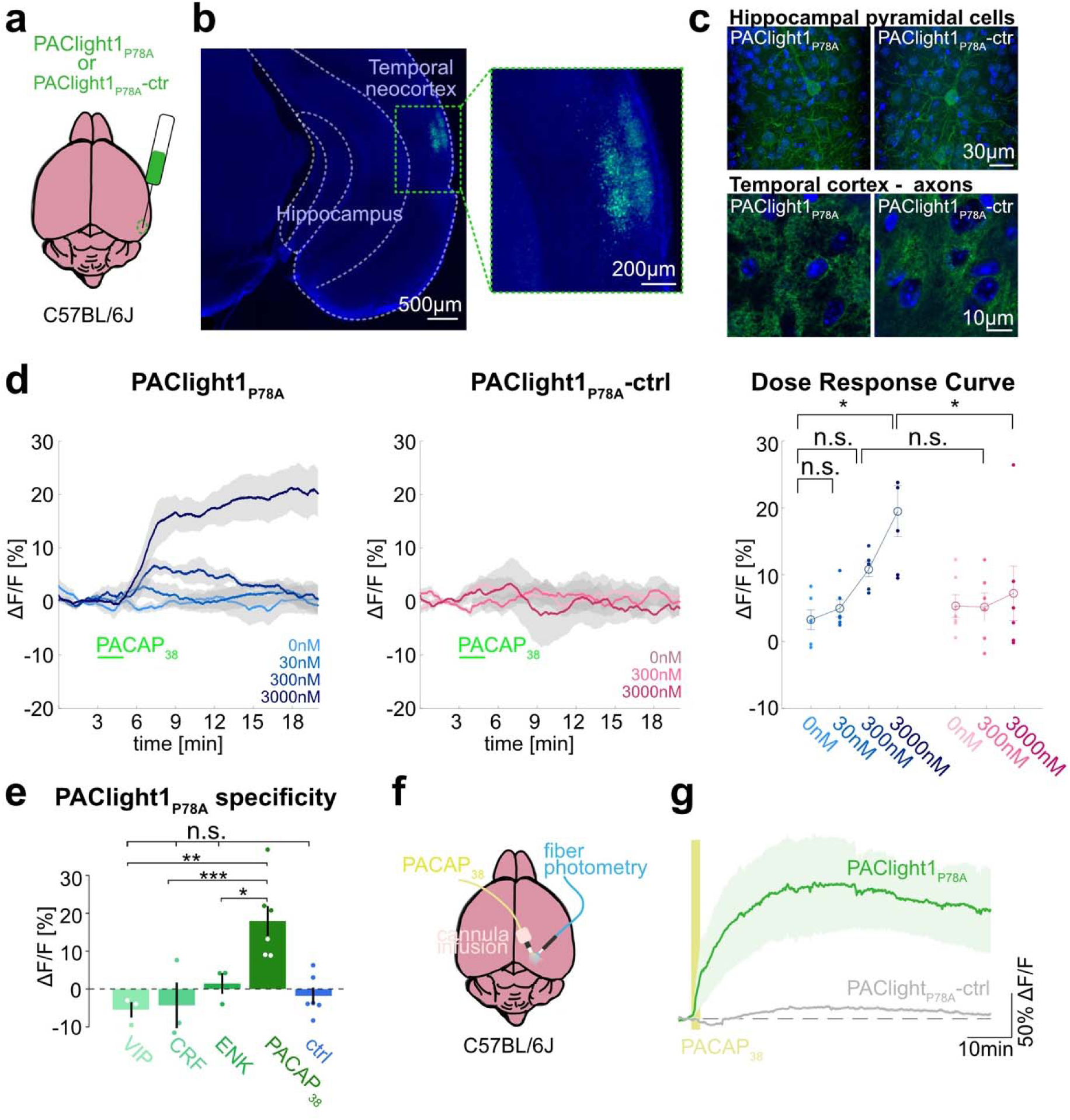
Ex vivo and in vivo sensor characterization in mice. **a)** AAVs encoding the PAClight1_P78A_ or PAClight1_P78A_-ctrl sensor were injected into the temporal neocortex of adult mice. **b)** Representative epifluorescent image of PAClight1_P78A_ fluorescence after 4 weeks of expression time. **c)** Maximum intensity projections of exemplary confocal images of PAClight1_P78A_ and PAClight1_P78A_-ctrl-expressing neurons and neuropil in hippocampus and cortex enhanced with GFP-immunostaining and counterstained with DAPI (blue). **d)** Acute mouse brain slices expressing PAClight1_P78A_ (left) and PAClight1_P78A_-ctrl (middle) were used to test the sensitivity of the sensor in the mammalian brain. PACAP_1-38_ was bath-applied for 2min (green bar) at indicated concentrations. Data shown as mean ± standard error of the mean (SEM). *N*=6 slices per condition from ≥3 mice. Right: Dose-response curve for PAClight1_P78A_ ΔF/F_0_ (blue shades) and PAClight1_P78A_-ctrl ΔF/F_0_ (pink shades) in response to indicated concentrations of PACAP_1-38_. Mann-Whitney-U-tests with Bonferroni corrections revealed no statistically significant difference in mean PAClight1_P78A_ peak responses to bath-application of 30nM (*p*=2.909), and 300nM PACAP_1-38_ (*p*=0.0519) compared to 0nM PACAP_1-38_. We found a significant difference between 0nM and 3000nM PACAP_1-38_ (*p*=0.013). Comparing mean peak ΔF/F_0_ PAClight1_P78A_ responses with PAClight1_P78A_-ctrl revealed a significant difference when bath-applying 3000nM (*p*=0.041), but not for 300nM (*p*=0.084). **e)** Quantification of mean PAClight1_P78A_ peak responses to bath-application of 3µM VIP, CRF, ENK, PACAP_1-38_ and negative control (ctrl) in acute mouse brain slices. *N*=3-6 slices per condition from ≥3 mice. No statistically significant difference (*F*_(4,10)_=5.15) was found between ctrl and VIP (*p*=0.9705), ctrl and CRF (*p*=0.9937) and ctrl and ENK (*p*=0.9726). A statistically significant difference (*F_(3,8)_=*9.19) was detected between PACAP_1-38_ and VIP (*p*=0.007), PACAP_1-38_ and CRF (*p*=0.001) and PACAP_1-38_ and ENK (*p*=0.045). Statistically significant differences are indicated with an asterisk, non-significant differences with n.s. **f)** AAVs encoding the PAClight1_P78A_ or PAClight1_P78A_-ctrl sensor were injected into the neocortex of adult mice. Fiberoptic cannulae and acute microinfusion cannulae were implanted nearby. **g)** PAClight1_P78A_ and PAClight1_P78A_-ctrl fluorescence changes upon microinfusion of 300µM PACAP_1-38_ (200nl) recorded with fiber photometry in freely behaving mice. *N*=5 PAClight1 _P78A_ and 4 PAClight1 _P78A_-ctrl mice. Data shown as mean ± SEM.

To investigate the sensor response dynamics and its sensitivity, acute brain slices expressing PAClight1_P78A_ or PAClight1_P78A_-ctrl were prepared and PACAP_1-38_ was bath-applied at concentrations ranging from 0nM to 3000nM. We observed a dose-dependent fluorescence increase to PACAP_1-38_ using PAClight1_P78A_, but not PAClight1_P78A_-ctrl (**Figure 4d**). PAClight1_P78A_ fluorescence increased by +6.8 and +17.5% ΔF/F_0_ when bath-applying 300 and 3000nM of PACAP_1-38_ respectively (**Figure 4d**). Comparing PAClight1_P78A_ fluorescence with PAClight1_P78A_-ctrl fluorescence revealed a statistically significant difference when bath-applying 3000nM of PACAP_1-38_ (**Figure 4d**).

To assess the specificity of PAClight1_P78A_ to PACAP_1-38_ in comparison to other neuropeptides in acute brain slices, we recorded its response to 3µM VIP, corticotropin-releasing factor (CRF), and enkephalin (ENK). PAClight1_P78A_ clearly increased its fluorescence in response to 3µM PACAP_1-38_ (+17.5% peak ΔF/F_0_), but not to VIP (−5.5% peak ΔF/F_0_), CRF (−4.3% peak ΔF/F_0_), or ENK (+1.4% peak ΔF/F_0_) (**Figure 4e**).

In conclusion, our data show that in acute mouse brain slices, PAClight1_P78A_ detects concentrations of >300nM of PACAP_1-38_ in superfused bath application, while not reacting to other neuropeptides tested.

### In vivo PACAP detection in behaving mice

To test whether PAClight1_P78A_ can be used to characterize ligand binding and diffusion in vivo in behaving mice, we implanted fiberoptic cannula into the neocortex of mice expressing PAClight1_P78A_ or PAClight1_P78A_-ctrl to image in vivo fluorescence dynamics while microinjecting PACAP_1-38_ through nearby cannula (**Figure 4f**). Microinfusion of PACAP_1-38_ (300µM, 200nl) led to peak fluorescence increases of 165.5 ± 58.0% ΔF/F_0_ in PAClight1_P78A_-expressing mice. The fluorescence peaked at 28 min and dropped to 128.6 ± 45.7% ΔF/F_0_ 1 hour after, when injections were positioned in average 318µm from the recording site. In contrast, PAClight1_P78A_-ctrl-expressing mice showed a fluorescence increase of only 16.7 ± 2.6% ΔF/F_0_ (**Figure 4g**).

In conclusion, we show that PAClight1_P78A_ can be a useful tool to detect PACAP_1-38_ in vivo with a dynamic range that allows for the detection of drug injections even in the presence of intact endogenous PACAP systems in mice. Moreover, our data suggest that PACAP_1-38_ diffuses efficiently across hundreds of µm, with slow extracellular degradation in neocortical brain areas of mice.

### Two-photon validation of the sensor in living zebrafish

Zebrafish larvae are an important animal model that has long been recognized for its utility and applicability to drug discovery^32–34^. Moreover, PAC1 signaling has been associated with the adaptive stress response of zebrafish^9,35^. The high degree of amino acid sequence conservation of PACAP across the phylogenetic tree, as well as the *in vitro* response of PAClight1_P78A_ to zebrafish PACAP that we observed, motivated us to also functionally validate our sensor for use in live zebrafish larvae. Based on a publicly available single-cell RNA sequencing (scRNAseq) dataset^36^, we identified the olfactory region of 4 day post-fertilization (dpf) old zebrafish larvae to express high levels of *Adcyap1b* in ∼75% of cells composing the olfactory region. The presence of *Adcyap1b* expression in the olfactory region was further confirmed with data obtained by Fransworth et al.^37^ (**Figure 5a**). To induce expression of PAClight1_P78A_ in the olfactory region, we used the gal4 driver line Tg(GnRH3:gal4ff), which can strongly drive expression (e.g. of GCaMP6s) in the olfactory bulbs (**Figure 5b**). In conjunction with this gal4 driver line, we used Tol2-mediated integration of a UAS-promoted PAClight1_P78A_ construct. At 4dpf we immobilized the larvae with low-melting point agarose and performed two-photon volumetric imaging of the olfactory region for 5 consecutive 3D volumes as baseline. The immobilized zebrafish larvae were then placed onto a micromanipulator platform to inject 50nL of a 1mM PACAP_1-38_ in saline solution or 50 nL of saline only (negative control) into the ventricular space (i.e. ICV injection). Subsequently, the larvae were re-imaged for 15 additional 3D volumes to record potential alterations in the pixel intensity values emitted by the PAClight1_P78A_ sensor (**Figure 5c, 5c’**). We observed a strong increase in PAClight1_P78A_ fluorescence already at ∼120 seconds post-ICV injection with ΔF/F_0_ levels continuously rising until the end of the post-ICV injection recordings (**Figure 5d, 5d’’**). Peak ΔF/F_0_ levels and the Area Under the Curve significantly differed between PACAP_1-38_/saline and saline-only injected animals (n_saline_= 5, n_PACAP1-38_= 5, Kolmogorov-Smirnov test *p*=0.0079, **Figure 5e, 5e’**). The condition median for peak ΔF/F_0_ levels was 70-fold larger in PACAP_1-38_/saline injected animals (145.76% ΔF/F_0_) than in saline-only injected animals (2.06% ΔF/F_0_) (**Figure 5e**). However, we noticed considerable variability in the extent of the dynamic range displayed by different ROIs within the same and also between different zebrafishes (**Figure 5d’’, 5e’**).

**Figure 5.**
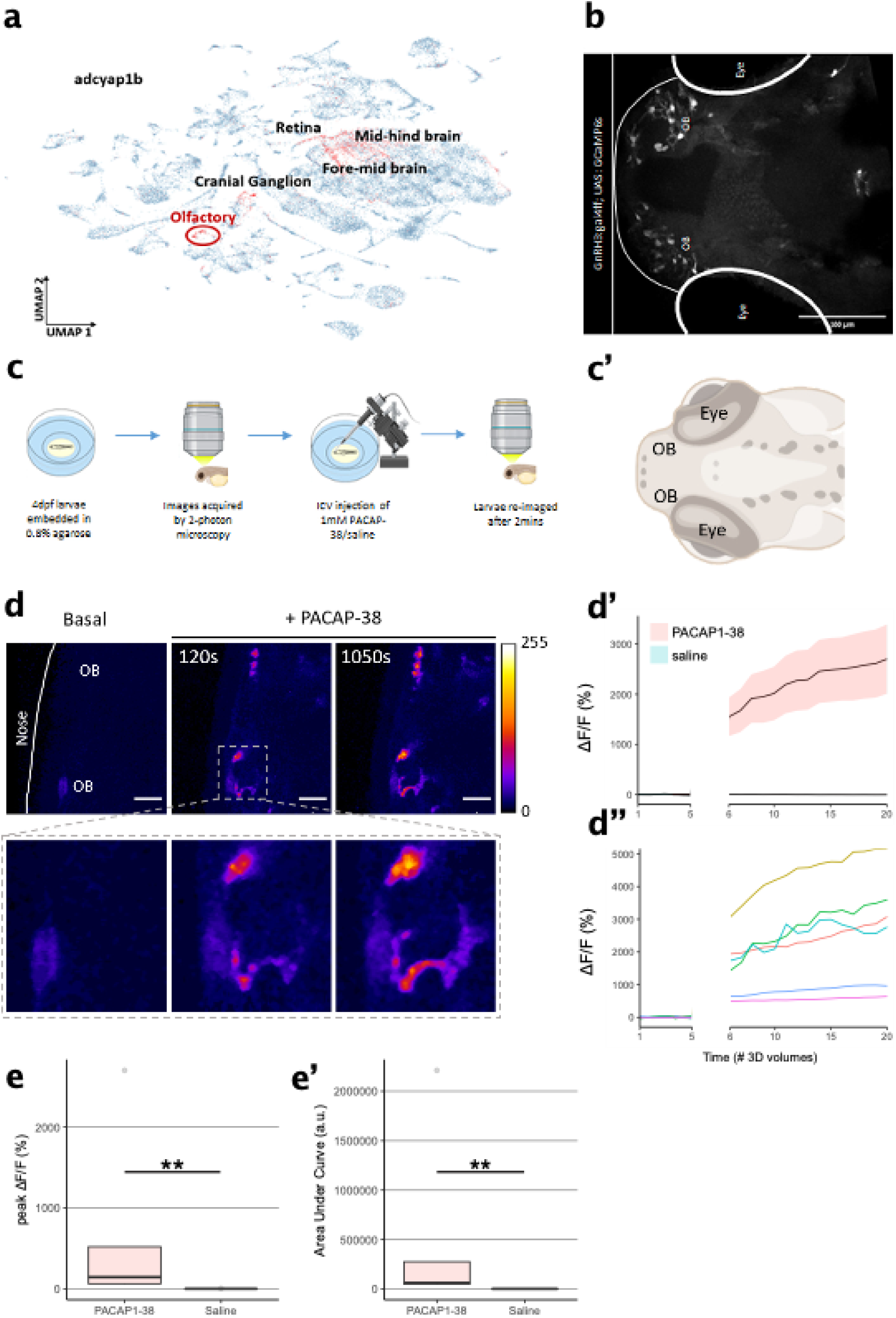
Characterization of PAClight1 in live zebrafish larvae. **a)** Uniform manifold approximation and projection (UMAP) showing topological distribution of single cell gene expression clusters with high adcyap1b (PACAP) expression of 0-4 dpf larvae. High adcyap1b expression is highlighted in the olfactory bulb. **b)** Maximum intensity projection image of the 4dpf *Tg(GnRH3:gal4ff; UAS:GCaMP6s)* larvae shows GnRH3 expressing cells in the olfactory bulb. **c)** Schema representing the experimental design. 4dpf larvae were immobilized and 3D volumetric images across time were obtained during the naïve state. 50nl of 1mM PACAP-38/saline was injected intracerebroventricularly and the same larvae was imaged again after 2mins. **c’)** Schema of imaged region of interest including the olfactory bulb in the 4dpf larvae. **d)** (top) Representative maximum intensity projection of two-photon volumetric images showing naïve sensor at baseline and its increase in fluorescence after PACAP-38 injection in the same larvae along time (scale bar = 20um). Color bar is representative of the ΔF/F_0_ in the images. (bottom) Zoomed-in image of the left olfactory bulb showing increase in fluorescence after injection of PACAP_1-38_. **d’)** Representative quantification of the change in fluorescence with respect to basal fluorescence depicted as ΔF/F_0_. PACAP_1-38_injected larvae show an increase in activity of the fluorescent sensor as compared to their saline injected sibling controls. **d”)** Individual cell traces show variability in sensor responses in different cells that may be a function of expression of the sensor on the cell surface. **e,e’)** Peak ΔF/F_0_ and Area under the Curve (AUC) of PACAP_1-38_ injected larvae is significantly higher than the saline injected controls (n_control_ =5, n_PACAP-38_ =5, Kolmogorov-Smirnov test *p*=0.0079)

## Discussion

In this work we engineered a new family of genetically-encoded fluorescent sensors using the human PAC1R as a GPCR scaffold. Two of the sensors that we developed (PAClight1 and PAClight1_P78A_) exhibit a very high dynamic range (above 1000% ΔF/F_0_), excellent expression at the cell surface, high basal brightness, and retain the pharmacological profile and ligand-binding profile of the parent receptor, with the exception of PAClight1_P78A_ whose response to VIP is intentionally abolished via a single point mutation. Given the very high sensitivity of these tools, future work could explore whether a grafting-based approach, similar to the one we recently described for class-A GPCR-based sensors^18^, could lead to the direct generation of multiple class-B1 sensors based on the optimized fluorescent protein module from PAClight.

As part of this sensor family, we also introduced a control sensor harboring the two point mutations R199A and Y161A, in which the response to PACAP_1-38_ is abolished. Given the higher PACAP-selectivity of PAClight1_P78A_, this sensor variant is intended to be employed when there is an experimental need to ensure maximal selectivity of the response for the endogenous ligand PACAP_1-38_ (for example if the sensor is to be used for attempting the detection of endogenous PACAP_1-38_ release). For all other experimental scenarios, PAClight1 can be a better choice, as it retains the original wild-type sequence in its ECD and thus most closely represents the natural ligand-binding profile of the human PAC1 receptor.

Importantly, we demonstrate successful expression and functionality of PAClight1_P78A_ and PAClight1_P78A_-ctrl in the mouse brain by injecting AAVs encoding the sensor stereotactically. Additionally, our experiments in zebrafishes and mice revealed intriguing inter-and intraindividual differences in maximal sensor response to ICV and intracranial injections of the peptide ligand. These differences could potentially stem from variances in the expression density of PAClight1_P78A_ within single cells. An intriguing alternative explanation, supported by the observation of continuously increasing ΔF/F_0_ levels (at least within the first 15 min), is that variations in diffusion rates and spatial diffusion patterns of PACAP_1-38_ within the brain parenchyma and cortical tissue may contribute to these differences. Furthermore, the distance between PAClight1_P78A_ -expressing cells and the site of injection might influence the extent of fluorescent response. Under more controlled experimental conditions, PAClight1/PAClight1_P78A_ - expressing zebrafishes and mice could therefore potentially be used to investigate and validate diffusion properties of PAC1R ligands in live organisms at a relatively high-throughput or to screen for novel small-molecule or peptide PAC1R agonists and antagonists with improved penetration properties across the ventricular wall.

Consistent with our findings in zebrafish, PACAP_1-38_ microinjections into the mouse cortex revealed slow increases in ΔF/F_0_ levels even when PACAP_1-38_ was injected only 100s of µm away from the recording site, substantiating the hypothesis of slow diffusion of PACAP_1-38_ through brain tissue. Furthermore, our results show a sustained activity with a slow decay of PAClight_P78A_ fluorescence, especially at high concentrations. We hypothesize that this delayed decay may result from the slow clearance of peptides from extracellular space, putatively due to saturation of the enzymatic degradation system. An alternative explanation could be a long-lasting activation of PAC1 after binding of PACAP_1-38_. This explanation is substantiated by slow decay in our slice experiments and by previous findings that PACAP_1-38_ effects on downstream PKA signaling is longer-lasting than effects of two other tested peptides (VIP and CRH)^38^.

Notably, in this work we did not demonstrate the use of any of our PAClight sensors for the detection of endogenous PACAP release in tissues or animal models, as our focus was deliberately set on the characterization and validation of the tools for applications to drug screening and/or development across species. Future work could focus on employing these tools to test whether or not they could be suitable for detecting endogenous PACAP dynamics with high spatiotemporal resolution in tissues or awake behaving animals. Based on our observations from mouse slice and in vivo recordings, bulk or epifluorescent imaging is unlikely to reveal endogenous release of PACAP, given that measured peptide concentrations typically occur in the femtomolar to nanomolar range^39^. However, further optimization of the sensor to increase sensitivity and brightness coupled with the use of two-photon imaging that allows to focus on PAClight-expressing cell membranes nearby PACAP release sites, holds promise for future application of this sensor for in vivo detection of endogenously released PACAP. Considering the slow diffusion of PACAP_1-38_ observed in our in vivo experiments, it will be quintessential to record PACAP release close to the main sites of action to capture physiologically relevant temporal dynamics. This could be achieved for example also by expressing the sensor exclusively in cells that express PACAP-sensitive receptors (PAC1, VPAC1, VPAC2) and then trigger PACAP release via optogenetic stimulation or behavioral paradigms.

Our results show that the new tools introduced in this study can be valuable assets to investigate real-time dynamics of PAC1 receptor activation in response to the application of specific PAC1 receptor agents, both at a cellular level and in vivo in zebrafish and in the mammalian brain. The in vivo validation of our sensor demonstrates its ability to reveal diffusion dynamics of applied drugs and peptides within brain tissue. We therefore suggest that this tool can be employed in applications such as studying the diffusion of novel PAC1-targeting drugs and peptides, thereby offering mechanistic insights into the localization of drug actions in the brain.

## Methods

### Molecular cloning

A synthetic DNA geneblock for the human PAC1R-null receptor sequence (PAC1R) was designed and ordered (Life Technologies) based on the NCBI protein data bank entry “NP_001109.2”. The protein-coding sequence was codon optimized and flanked by HindIII (5’) and NotI (3’) restriction sites for cloning into a pCMV plasmid (Addgene #60360). A hemagglutinin (HA) signal peptide (MKTIIALSYIFCLVFA) was introduced 5’ to the PAC1R coding sequence. Circular Polymerase Extension Cloning (CPEC) was used to replace the intracellular loop 3 (Q336-G342) of PAC1R with a cpGFP module from dLight1^10^. For sensor optimization, libraries of sensor variants were created through site-directed mutagenesis. For signaling assays, a linker + SmBit sequence^40^ (GNSGSSGGGGSGGGGSSGG + VTGWRLCERILA) was cloned at the 3’end of both the PAClight1_P78A_ and the PAC1R sequence. The XhoI cleavage site from within the SmBit linker sequence and the first 4 residues of the SmBit linker (GNSG) were first cloned into the 3’end of the PAClight1_P78A_ and PAC1R sequences using PCR. Subsequently, a restriction digest using XhoI and XbaI restriction enzymes was performed on the SmBit containing plasmid (B2AR-SmBit) as well as on the pCMV_PAClight1_P78A_ and pCMV_PAC1R plasmids. Ligation of the linker+SmBit insert into the linearized backbones was performed after gel extraction and purification of the insert. For generation of stably expressing and inducible Flp-In T-REx 293 cells, the PAClight1_P78A_ and PAClight1 sequences were cloned into the pcDNA5/FRT/TO backbone, respectively, using restriction digest with BamHI and NotI restriction enzymes. PCR reactions were performed using a Pfu-Ultra II Fusion High Fidelity DNA Polymerase (Agilent), whereas Gibson Assembly was performed using NEBuilder HiFi DNA Assembly Master Mix (New England Biolabs). All sequences were verified using Sanger sequencing (Microsynth).

### Structural modelling, protein sequence alignment and peptide synthesis

Modelling of protein structure for the PAClight1_P78A_ sensor construct was performed using AlphaFold2^41^, using pdb70 as a template mode. The best-scoring prediction was then manually edited using UCSF Chimera (version 1.13.1). Multiple sequence alignment was performed using ClustalOmega^42^ and visualized with the Jalview^43^ software (version 2). Zebrafish PACAP2_1-27_(zfPACAP2), chicken PACAP_1-38_ (chPACAP38), amnesiac, maxadilan (Maxadilan_1-61_), and Max.d.4 peptides were synthesized on an automated fast-flow peptide synthesizer (AFPS) using a previously described protocol^44^.

### Cell culture, imaging and quantification

HEK293T cells (ATCC #CRL-3216) were cultured in DMEM medium (Thermo Fisher) supplemented with 10% FBS (Thermo Fisher) and 1x Antibiotic-Antimycotic (100 units/mL of penicillin, 100µg/mL of streptomycin, and 0.25 µg/mL of Amphotericin B, Thermo Fisher) and incubated at 37°C with 5% CO_2_. Cells were transfected at 50-60% confluency in glass-bottomed dishes using the Effectene transfection kit (Qiagen) according to manufacturer instructions, and imaged 24-48h after transfection, as previously described^16^. Primary cultured hippocampal neurons were prepared and transduced as previously described^18^. Before imaging, all cells were rinsed with 1mL of Hank’s Balanced Salt Solution (HBSS, Life Technologies) supplemented with Ca^2+^ (2mM) and Mg^2+^ (1mM). Time-lapse imaging was performed at room temperature (22° C) on an inverted Zeiss LSM 800 confocal microscope using either a 40x oil-based or a 20x air objective. Longer-term imaging for determination of internalization/stability as well as reversibility was performed using a 10x air objective, with 1 frame (1024×1024 pixels) acquired every minute (2x pixel averaging). Imaging was performed using a 488nm laser as excitation light source for PAClight1 sensors. During imaging, ligands were added in bolus on the cells using a micropipette to reach the final specified concentrations of ligands on the cells. For quantification of recordings in single dishes, an average intensity projection across the whole temporal stack was first generated. Pixel-intensity based thresholding in Fiji was then performed manually on the average intensity projection to segment the plasma membrane as accurately as possible. Using the magic wand tool from Fiji, connected patches of segmented plasma membranes (sampled from multiple cells) were selected as regions of interest (ROIs). The ROIs were then projected onto the temporal stack to make sure the plasma membrane did not drift out of the ROI during the time lapse recording. Sensor response (ΔF/F_0_) was calculated as the following: (F(t)-F_base)/F_base with F(t) being the ROI fluorescence value at each time point (t), and F_base being the mean fluorescence of the 10 time points prior to ligand addition.

### Stable cell line generation and maintenance

The stable cell line for tetracycline-inducible expression of PAClight1_P78A_ was generated following previously described procedures^27^. Flp-In T-REx 293 cells were grown in DMEM supplemented with 10% (v/v) fetal bovine serum (FBS; Invitrogen), 100µg/mL zeocin (ThermoFisher Scientific), and 15µg/mL blasticidin (ThermoFisher Scientific). To obtain a cell line with stably-integrated PAClight1_P78A_ or PAClight1 expression cassettes, cells were grown in T150 flasks (Corning) until 70% confluency, were then transfected with 0.6µg of pcDNA5/FRT/TO-PAClight1_P78A_ DNA vector and 5.4µg pOG44 vector using Effectene transfection kit (Qiagen). Two days after transfection, cells were split and the medium was changed to DMEM supplemented with 10% (v/v) FBS, 200µg/mL Hygromycin B (Sigma), and 15µg/mL blasticidin. The medium was then replaced twice a week until individual colonies were visible. An individual colony was manually selected and expanded for subsequent experiments. Induction of PAClight1_P78A_ or PAClight1 expression was obtained by adding 1µg/mL doxycycline (Sigma) to the cell medium 1-2 days prior to experimentation.

### Spectral characterization of the sensors

One-photon spectral characterization of the PAClight1_P78A_ sensor was performed using PAClight1_P78A_-transfected HEK293T cells before and after addition of PACAP_1-38_ (10µM). One-photon fluorescence excitation (λ_em_ = 560nm) and emission (λ_exc_ = 470nm) spectra were determined on a Tecan M200 Pro plate reader at 37°C. 24 hours after cell transfection in 6-well format (linear PEI), ∼1Lmillion cells were dissociated with addition of TrypLE Express (ThermoFisher) and thoroughly washed with PBS. Next, cells were resuspended in 300µl of PBS and aliquoted into two individual wells of a 96-well microplate with or without PACAP_1-38_ (10µM), together with two wells containing the same amount of non-transfected cells to account for autofluorescence and a single well containing PBS for subtraction of the Raman bands of the solvent.

### Flow cytometry

HEK293T cells and stable HEK293_PAClight1_P78A_/PAClight1 cells were seeded into T175 flasks and grown to 50-60% confluency under culture conditions described above. HEK293T cells were then transfected with 20µg of pCMV_PAClight1_P78A_ or pCMV_PAClight1 using linear PEI (Sigma-Aldrich; #764965) and a PEI-to-plasmid ratio of 3:1. Stable HEK293_PAClight1_P78A_/PAClight1 cells were induced with 1µg/mL doxycycline (Sigma) at 50-60% confluency. Two days after transfection or induction, cells were washed in 1x PBS before detachment with TrypLE Express (ThermoFisher) and subsequent centrifugation at 150xG for 3 minutes. Palleted cells were then resuspended in ice-cold FACS buffer containing 1xPBS, 1mM EDTA, 25mM HEPES (pH 7.0), 1% fetal bovine serum and diluted to 4-4.8×10^6^ cells/mL. A dilution series of PACAP_1-8_ and VIP was prepared in FACS buffer. 125µL of peptide solution was pipetted into a 96-well format and another 125ul of cell suspension was added to each well. The 96-well U-bottom plate was then loaded into a BD FACS Canto II cytometer equipped with a high-throughput sampler for 96-well format sampling. Voltage for the photomultiplier tubes was set to 200V for the forward scatter detector, to 400V for the side scatter detector, and to 350V for the FITC channel fluorescence detector. Excitation was performed at 488nm, while emission was directed through a 502LP mirror and a 530/30 band pass filter. Sampling from 96-well format wells was performed with an initial mixing step (3× 100µl mixes at 150µL/s) followed by sample acquisition of 200µL at a flow rate of 1-3µL/s. Sampling was performed until 100K events were recorded per well. After sample acquisition of each well, a 800µL wash step of the sampling line was performed. Sample acquisition within a dilution series replicate was performed from low-to-high concentrations to minimize potential peptide carry-over into the neighboring wells. Furthermore, a minimum of six wells without PACAP or VIP present were sampled between each replicate of the same construct. Raw data was exported in FCS3.1 format and further processed for analysis within R. The following R packages were used for the analysis and visualization of the flow cytometry experiments: flowCore, flowAI, ggcyto, dplyr, tidyr, forcats, purrr, stringr, openCyto, openxlsx, svglite, ggprism, ggnewscale, and glue. After import of the raw datasets, a rectangular gate (“Cells”) was defined on a FSC-A vs. SSC-A scatter plot with the limits being set to 25K-infinity and 15K-infinity, respectively. Next, a “Singlet” gate within the parent gate “Cells” was created automatically using the openCyto::singletGate function based on the FSC-A vs. FSC-H scatter plot. Median fluorescence intensities (MFI) of the FITC-A channel for all recorded events within the “Singlet” gate were calculated for each well. The MFI values were then normalized (“NormFITC_A”) within each titration replicate to the average of the 0nM peptide condition. Dose-response-curves and EC_50_ values were obtained by first grouping datasets by construct and peptide conditions. For each group, a non-linear least squares model was fit using the following formula:

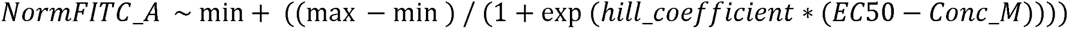

For the VIP titration on the non-responsive sensor variant, a linear regression model was fit using the following formula:

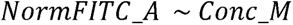

### NanoLuciferase complementation assays

HEK293 cells were seeded in 6-well plates and transfected with 0.25μg LgBiT-miniG (miniGs or miniGsq^45^) or LgBiT-β-arrestin2^40^ and 0.25μg SmBiT-tagged receptor plasmids using 3µL Lipofectamine 2000 (Thermo Fisher). 24 hours after transfection, cells were plated onto black clear-bottomed 96-well plates at 50,000 to 100,000 cells/well or 384-well plates at 20,000 to 30,000 cells/well in DMEM without phenol red, supplemented with 30mM HEPES (pH 7.4) and Nano-Glo Live Cell Reagent (Promega), and incubated for 45 min at 37°C. Luminescence signal was measured simultaneously across the plate using the FDSS/μCELL plate reader (Hamamatsu). Baseline luminescence (before agonist addition) was acquired for 3min. Vehicle (buffer) or agonist (PACAP_1-38_, 1μM) were added simultaneously into wells by an integrated dispensing unit. Luminescence was recorded every 1 – 2 s for 8 min post agonist addition. Agonist-treated wells were initially normalized to vehicle wells, luminescence intensity was then normalized to the baseline prior to ligand addition.

### Generation and imaging of transgenic fishes

The PAClight1_P78A_ sequence was cloned into a Tol2 transposase-based plasmid^46^ with a UAS promoter. The plasmid (75ng/ul) was injected into *Tg(GnRH3:gal4ff)*^47^ embryos at the single-cell stage along with transposase mRNA (25 ng/ul). Embryos were sorted at 3 days post fertilization using the red heart marker (cmlc:mCherry) in the *UAS:PAClight1_P78A_*construct. Positive zebrafish larvae were immobilized using alpha-bungarotoxin (1mg/ml for 30 s) and mounted ventrally (ventral side down) on to a custom molded 3 cm plate using 0.8% low-melt agarose (Sigma-Aldrich). Plates were perfused with E3 media (5mM NaCl, 0.17mM KCl, 0.33mM CaCl_2_, 0.33mM MgSO_4_ in H_2_O) prior to imaging. Larvae were imaged under the Leica TCS SP8 Multi-Photon Microscope with a 25X/0.95NA water immersion objective. Volumetric images across time were obtained after irradiation at 890 nm at 400 Hz/frame with 2x averaging. Larvae were imaged for five volumes in the naïve state followed by intracerebroventricular injections (ICV) of 1mM PACAP-38 (Sigma-Aldrich) in saline solution (0.6% NaCl, 0.02% Na_2_CO_3_) and only saline for controls. Larvae were then re-imaged approximately 120 seconds after ICV injection using the same imaging parameters for 15 volumes. Basal and post-injection images were concatenated and registered using descriptor-based series registration on ImageJ/Fiji (https://github.com/fiji/Descriptor_based_registration.git). Raw fluorescence traces were extracted from these images from manually drawn ROIs around cells. Statistical analysis and plotting of graphs were performed in MATLAB and R. Baseline fluorescence, F_0_, was computed using basal fluorescence intensities, which were then used to calculate ΔF/F_0_. The Kolgomorov-Smirnov t-test was performed where applicable. Single cell transcriptomic data^37^ from 4dpf zebrafish larvae was plotted as a function of UMAPs in the UCSC cell browser. High expression of adcyap1b (PACAP) was visualized in Cluster 94 (https://cells.ucsc.edu/?ds=zebrafish-dev&gene=ENSDARG00000027740). Tg(GnRH3:gal4ff) transgenic line was used to express the PAClight1_P78A_ in the olfactory bulb (OB). To confirm that GnRH3 cells were indeed present in the OB at 4dpf, a UAS:GCaMP6s construct was expressed in the Tg(GnRH3:gal4ff) line.

### Virus production

The biosensor AAV constructs were cloned in the Patriarchi laboratory, while the opsin AAV construct was constructed by the Viral Vector Facility of the University of Zürich (VVF). All viral vectors were produced by the VVF. The viral titers of the viruses used in this study were: AAV9.hSyn.PAClight1_P78A_, 0.75-1.5 × 10^13^ GC/mL, AAV9.hSyn.PAClight1_P78A_-ctrl, 1.6 × 10^13^ GC/mL

### Animals

Rat embryos (E17) obtained from timed-pregnant Wistar rats (Envigo) were used for preparing primary hippocampal neuronal cultures. Wild-type C57BL/6JRj mice (Janvier, 6-10 weeks old) of both sexes were used in this study. Mice were kept in standard enriched cages with *ad libitum* access to chow and water on either normal or reversed 12-h/12-h light/dark cycle. Mice were housed in cages of 2-5 animals.

All procedures in mice were performed in accordance with the guidelines of Medical University of Vienna and under approved licenses by the Austrian Ministry of Science.

Zebrafish were maintained and bred by standard protocols and according to FELASA guide-lines. All experiments using zebrafish were approved by the Weizmann Institute’s Institutional Animal Care and Use Committee (Application number 06340722-1).

### Stereotactic surgeries

For ex vivo slice imaging and validation of mouse brain expression, 6 to 10-week-old male and female C57BL/6JRj mice (Janvier) were anesthetized with 5% isoflurane and maintained under stable anesthesia during the surgery (1.3-2% isoflurane). Lidocaine (3.5mg/kg; Gebro Pharma, #100562 1404) and Carprofen (4mg/kg; Zoetis, #256684) were administered subcutaneously for local anesthesia and general analgesia. A small craniotomy was made bilaterally, targeting the temporal neocortex (Injection site at −4.1 mm posterior to bregma, as lateral as possible and 1.2 mm ventral to the pial surface). A glass micropipette was used for virus delivery and inserted slowly into the brain. Before and after virus delivery, the micropipette was kept in place for a waiting time of 8 min. 100-200nL of AAV9.hSyn.PAClight1_P78A_ or AAV9.hSyn.PAClight1_P78A_-ctrl were injected, with an injection rate of 40nL/min using a microsyringe pump (KD Scientific; #788110). The pipette was withdrawn slowly and the skin of the skull was sutured. The mice were allowed to recover for 4 weeks before being used for the experiments, with Carprofen (4 mg/kg) being administered on the two consecutive days after the surgery.

### Slice imaging

The mice were deeply anesthetized with isoflurane before being sacrificed by a transcardial perfusion with 30mL cooled sucrose solution (212mM sucrose, 3mM KCl, 1.25mM Na2H2PO4, 26mM NaHCO3, 1mM MgCl2, 0.2mM CaCl2, 10mM glucose) oxygenated with carbogen gas. The mice were decapitated and the brain was dissected out. 300µM coronal slices were prepared using a Leica VT1200 vibratome in cold oxygenated sucrose solution (0.12 mm/s, 0.8 mm amplitude). The slices were incubated in oxygenated Ringer’s solution (125mM NaCl, 25mM NaHCO3, 1.25mM NaH2PO4, 2.5mM KCl, 2mM CaCl2, 1mM MgCl2, 25mM glucose) at 37°C for 10min and afterwards kept at room temperature. The slices were imaged using an Olympus BX51WI microscope with an ORCA-Fusion Digital CMOS camera (Hamamatsu, #C14440-20UP) and a 10x objective (UMPLFLN10XW objective, Olympus). An EGFP filterset was used to control emission and excitation spectra (470/40, 525/50; AFH #F46-002). Micromanager software ^48^ was used to synchronize the excitation LED with the camera shutter. Videos were taken at 4 Hz framerate using 100 ms exposure time per frame. Cortical areas with clearly visible baseline fluorescence were selected for imaging. Baseline recordings of 10-20 min were performed prior to data acquisition to allow the slices to accustom to the conditions. Bath temperature was set to ∼31°C. The experimental recording was 20 min long, with the respective peptide being infused from minute 3-5. The following peptides were used: PACAP_1-38_ (30-3000nM; Phoenix Pharmaceuticals; #052-05), VIP (3000nM; Phoenix Pharmaceuticals; #064-16), CRF (3000nM; Phoenix Pharmaceuticals; #019-06), and Met-Enkephalin (3000nM; Phoenix Pharmaceuticals; #024-35). Of note, since our preliminary data suggested a decrease in PAClight1_P78A_ fluorescence in response to 3 µM CRF, we enriched the Ringer’s solution with synaptic transmission and neuronal activity blockers (NBQX, CPP, CGP 55845, Gabazine, and TTX) to prevent recruitment of neuronal circuits that could lead to endogenous PACAP_1-38_ release (Tocris, biotechne; #Tocris 1262/50, #Tocris 0373/50, # Tocris0247/50, #Tocris 1248/50, #Tocris 1069/1).

Data analysis was performed using a custom-written Matlab script. Movement correction as well as bleaching correction was performed. The average fluorescence across the whole imaging window was normalized to the baseline fluorescence measured during the first 3 min of the recording prior to peptide infusion to calculate ΔF/F_0_: (F(t)-F_base)/F_base with F(t) being the fluorescence value at each time point (t), and F_0__base being the mean fluorescence of the first 3min. 6 slices from at least 3 different mice were averaged for calculating the dose-response curve. For PAClight1_P78A_ responses to VIP, CRF and ENK, 3 slices from 3 different mice were averaged.

### In vivo photometric imaging and microinfusion

Craniotomies were made as described for stereotactic injections. Tapered fiberoptic cannula implants (MFC_200/230-0.37_2mm_MF1.25_A45) with low autofluorescence epoxy were implanted into the right neocortex at 0.9mm depth with the angled (uncoated) side of the fiber tip facing towards the left. The implant was stabilized with cyanoacrylate glue.

Cannula were surgically implanted to locally infuse PACAP_1-38_ into the cortex. Stainless steel guide cannulae (26 gauge; C315GA/SPC, InVivo One, Roanoke, VA, USA) were positioned in cortical L1, such that the tip of the internal cannula terminated 400µm medial to an the fiberoptic cannula. The implant was secured with cyanoacrylate glue. Dummy cannulae that did not extend beyond the guide cannulae (C315DC/SPC, InVivo One) were inserted to prevent clogging.

Mice were handled for 7 days prior to drug infusions.

For photometric recordings, a 200µm diameter and 0.37 NA patchcord (MFP_200/220/900-0.37_5m_FCM-MF1.25, low autofluorescence epoxy, Dorics) was used to connect the implanted fiberoptic cannulae to a Dorics filter cube for blue (465-480nm) excitation light. Emission light was detected by built-in photodetectors (500-540nm). Signals from the photodetectors were amplified with Dorics built-in amplifiers and acquired using a Labjack (T7). LJM Library (2018 release) was used to allow communication between MATLAB and Labjack. The voltage output from the LED drivers was amplitude modulated at 171 Hz (sine wave) as described previously ^49^ Amplitude modulation was programmed in MATLAB. 470 nm LEDs (M470F3, Thorlabs; LED driver LEDD1B, Thorlabs) were used. Light power at the patchcord tip was set to an average of 45µW. Fluorescence was calculated with custom-written MATLAB scripts based on a previous publication (Owen and Kreitzer, 2019). Photometry data were sampled at 2052 Hz.

For drug infusions, dummy cannulae were replaced by internal cannulae (33 gauge; C315LI/SPC, InVivo One) that extended 1 mm beyond the guide cannulae and were connected to a microinfusion pump. After a recovery time of 5-10 min, PACAP_1-38_ (200nl; 300µM diluted in normal Ringer solution, NRR) was infused at 100 nl/min (NRR in mM: 135 NaCl, 5.4 KCl, 5 HEPES, 1.8 CaCl2, pH 7.2 adjusted with KOH, sterile-filtered with 0.2μm pore size).

### Histology

To verify expression of the sensor in the mouse cortex and hippocampus, virus injections were performed with stereotactic surgeries as described above. The virus was allowed to express for four weeks, before the mice were transcardially perfused with PBS and 4% paraformaldehyde, before the brain was dissected and processed into 50µm coronal slices using a Leica VT1000S vibratome. Slices were immunostained with chicken anti-GFP antibodies (Abcam Cat# ab13970, RRID:AB_300798; 1:500 diluted). In brief, slices were permeabilized and blocked for 1 h with PBS containing 5% NGS and 0.2% Triton X-100, followed by a 24 h antibody incubation at 4°C in PBS containing 5% NGS and 0.2% Triton X-100. Sections were washed in PBS and then incubated for 1 h in 1:500 diluted goat anti-chicken IgG (H+L) cross-adsorbed secondary antibody (Alexa fluor 488; Fisher Scientific A11039). The mounted slices were imaged using a Leica SP8 X confocal microscope equipped with a 63x 1.3 NA glycerol immersion objective.

### Statistical analysis

For pairwise analysis of sensor variants, the statistical significance of their responses was determined on a case-by-case basis using a two-tailed unpaired Student’s t-test with Welch’s correction. Analysis of variance (ANOVA) testing was followed by pairwise comparison with correction for multiple comparison with either Dunnett’s correction or Hochberg correction. For data not meeting assumptions of normality or homoscedasticity, Mann-Whitney-U tests followed by Bonferroni corrections were applied. All p-values are indicated either in the results section or in the figure legends. Data of sensor screening and validation experiments is displayed as mean +/−1 standard deviation. No statistical methods were used to predetermine sample size in cultured cells. Power calculations were performed to determine sample size for experiments using mice.

## AUTHOR CONTRIBUTIONS

TP led the study. TP and RBC conceived the development of the PAClight sensors. RBC performed all *in vitro* molecular cloning and screening that led to the development of PAClight sensors and characterized the optical and pharmacological properties of the sensor under the supervision of TP. AG measured and analyzed sensor kinetics under the supervision of TP. EW and PH synthesized peptides under supervision of NH. KA performed intracellular signaling assays under the supervision of MS. MAB prepared cultured neurons under the supervision of DB. SN and SM performed and analyzed imaging experiments in acute brain slices and in vivo photometry recordings and microinfusions under the supervision of SM. PR performed the in vivo experiments using zebrafish under the supervision of GL. TP and RBC wrote the manuscript with contributions from all authors.

## DATA AVAILABILITY

The DNA and protein sequence of the sensor developed herein have been deposited on NCBI (accession number OQ366523) and are available in Supplementary Data S1. Viral DNA plasmids have been deposited both on ADDGENE and on the UZH Viral Vector Facility (https://vvf.ethz.ch/). Viral vectors can either be obtained from the Patriarchi laboratory, the UZH Viral Vector Facility, or Addgene. Data presented in this manuscript are available from the authors upon reasonable request.

## CODE AVAILABILITY

Custom written code can be made available upon request.

## ACKNOWLEDGEMENTS

The results are part of a project that has received funding from the European Research Council (ERC) under the European Union’s Horizon 2020 research and innovation program (Grant agreement No. 891959) (TP). We also acknowledge funding from the Swiss National Science Foundation (Grant No. 310030_196455 and 310030L_212508), Olga Mayenfisch Foundation, and Hartmann Müller Foundation for Medical Research (TP). PR is supported by a research grant for student’s fellowship from the Benoziyo Endowment Fund for the Advancement of Science and by the Weizmann–CNRS Collaboration Program. GL lab is supported by the Israel Science Foundation (#349/21); Israel Ministry of Science and Technology (#3-16548) and Hedda, Alberto, and David Milman Baron Center for Research on the Development of Neural Networks. MS and KA are supported by a Swiss National Science Foundation Eccellenza Professorial Fellowship to M.S. (PCEFP3_181282). SM and SN have been funded by the Vienna Science and Technology Fund (WWTF) and the City of Vienna through project VRG21-015. We would like to thank Prof. Marco Celio for generously contributing financial support during the project.

---

**Extended Data Figure 1.**
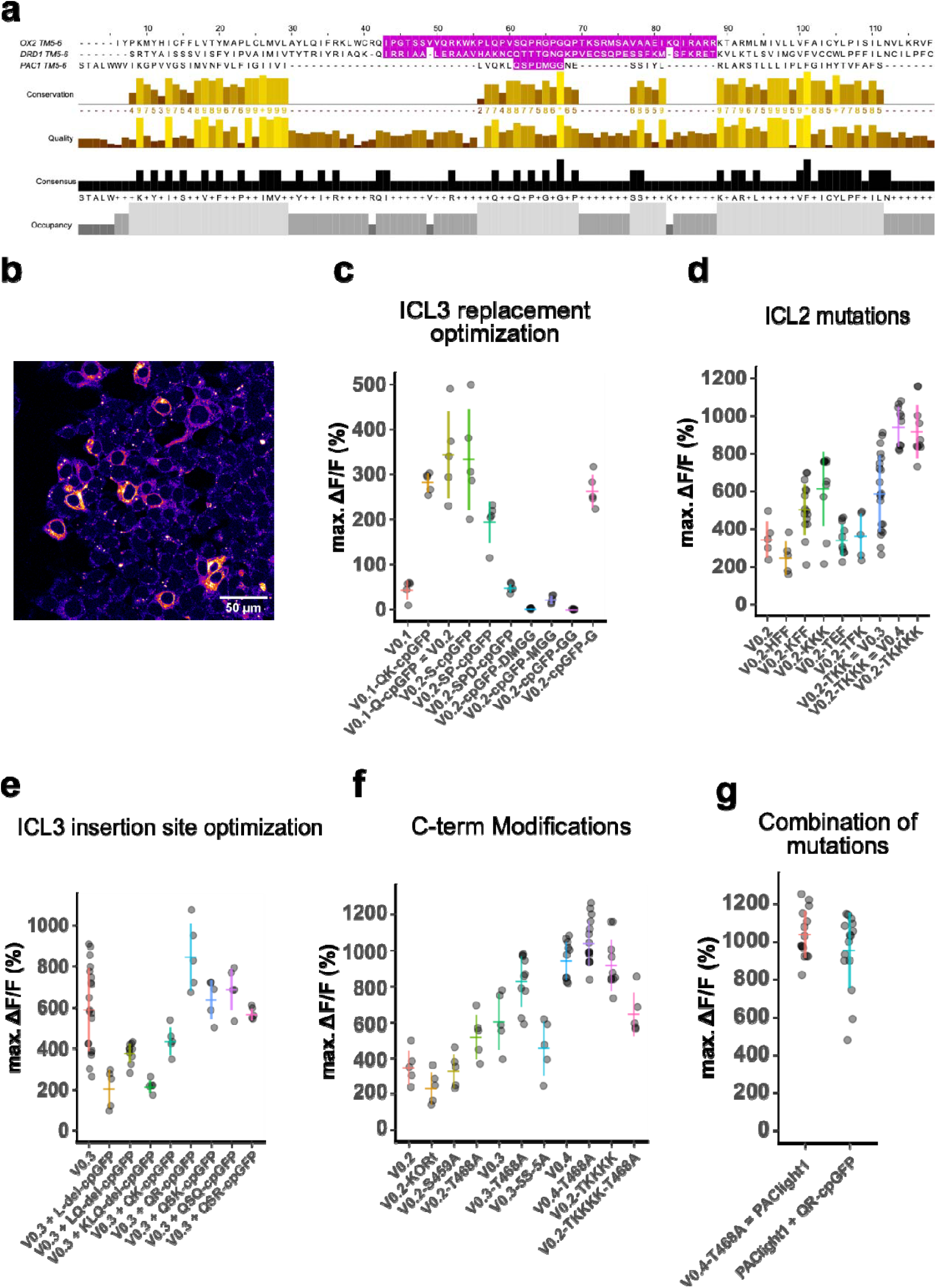
Optimizing the fluorescent response of PAClight sensors. **a)** Multiple amino acid sequence alignment of the TM5, ICL3, and TM6 domains of the human orexin 2 receptor (OX2), the human D1 dopamine receptor (DRD1), and of the human PAC1Rnull (PAC1). The sequences replaced by the cpGFP module in the OxLight1, dLight1.3b, and PAClight1_P78A_ sensors are highlighted in magenta. The OX2 and DRD1 receptor sequences align well due to the fact that they both belong to the class A GPCRs. The PAC1Rnull belongs to the class B1 family of GPCRs and hence aligns less well to the OX2 and DRD1 backbones. We therefore guided the initial design and site of cpGFP module insertion by protein structure rather than by amino acid sequence. **b)** Pseudo-colored confocal micrograph of HEK293T cells expressing the prototypical PAClight_V0.1. Plasma membrane expression can be detected, but a significant amount of sensor protein is retained intracellularly. **c)** Scatterplot representing insertion site variants screened for maximum dynamic range. Reinsertion of the glutamine on the N-terminal side of the cpGFP module (PAClight_V0.2) drastically increased the dynamic range to 343% ΔF/F_0_ (compared to 43.4% ΔF/F_0_ in PAClight_V0.1). **d)** Point mutations in the intracellular loop 3 (ICL3) further improved the dynamic range to 588% ΔF/F_0_ for PAClight_V0.3 (point mutations: F259K/F260K) and to 942% ΔF/F_0_ for PAClight_V0.4 (point mutations: F259K/F260K /P261K). **e)** On the backbone of the improved ICL2 variant PAClight_V0.3 we screened again for the optimal insertion site of the cpGFP module. To this end we performed serial deletions as well as insertions of charged residues on the residues n-terminal to the cpGFP module. This small-scale screen identified a mutant with a glutamine and arginine insertion before the cpGFP module (PAClight_V0.3+QR-cpGFP) with improved dynamic range (ΔF/F_0_ = 844%) compared to the PAClight_V0.3 backbone. **f)** To abolish β-Arrestin recruitment and further improve membrane localization, we further screened C-terminus mutants. We included mutants in this screen in which we replaced the C-terminus with the tail of the kappa opioid receptor (KORt), as well as point mutants of known post-translational modification sites and serine residues. The point mutation of the T468 to alanine further improved plasma membrane localization (data not shown) and improved the dynamic range from 942% ΔF/F_0_ (in PAClight_V0.4) to 1037% ΔF/F_0_. The resulting construct was named PAClight1. **g)** Since the insertion of QR in front of the cpGFP module had a positive effect on dynamic range on the backbone of PAClight_V0.3 **(e)**, we also tested this insertion of QR on the PAClight1 backbone. However, the QR insertion did not yield improved dynamic range on the PAClight1 backbone (ΔF/F_0_ = 953%) and was hence rejected. Each datapoint represents the average of the maxΔF/F_0_ within one ROI. In each replicate 5 ROIs each were quantified. *N*=4 for V0.3. *n*=3 for V0.2_KFF, PAClight1, and PAClight1+QR-cpGFP. *n*=2 for V0.2_KKK, V0.2_TEF, V0.2_KKKK, V0.3_LQ-del-cpGFP, V0.3_T468A, and V0.4. For all other constructs *n*=1 replicates each was screened. The y-axis position of the colored horizontal bars represents the mean across all replicates within a construct. The extent of the colored vertical bars represents +/−1 standard deviation. For ease of comparability some constructs are potted on multiple plots.

**Extended Data Figure 2.**
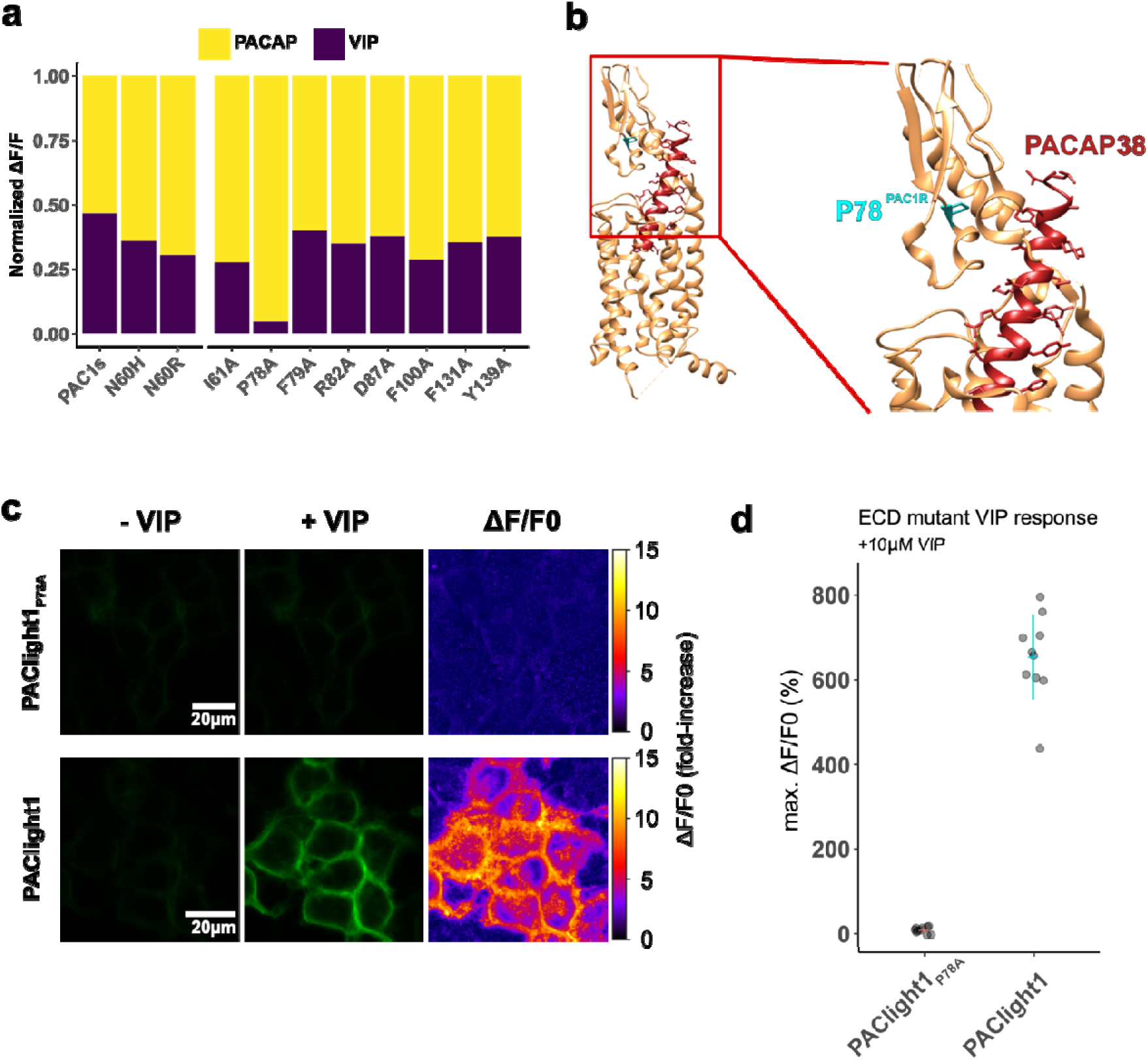
Engineering PACAP selectivity in PAClight1_P78A_. **a)** Stacked bar plot demonstrating the relative fluorescent response of ECD mutants towards 500nM of VIP followed by 9.09μM PACAP_1-38_. The mutant PAC1s contains the naturally occurring splice mutation (deletion) on the ECD. The N60H and N60R were inspired by the analysis of protein structure. The remaining alanine point mutants (right side of bar plot) were inspired by molecular dynamics (see main text). The P78A point mutation on the ECD resulted in excellent improvement of specificity. The highly PACAP-specific mutant was named PAClight1_P78A_. The data is normalized to the maximum response within each condition. *n*=1 for each mutant. **b)** Visualization of the ECD mutation P78A (cyan) on the backbone of a previously published cryo-EM structure of PAC1Rnull (beige) in interaction with PACAP_1-38_ (dark red) (PDB: 6LPB)^21^. **c)** Representative confocal micrographs of HEK293T cells expressing the PACAP-specific PAClight1_P78A_ and the VIP-responsive PAClight1 sensors before (left) and after (middle) 10μM VIP bath application. Pixel-wise calculation of ΔF/F_0_ is shown on the right. Note the complete absence of a fluorescent response to VIP in the PAClight1_P78A_ mutant. **d)** Quantification of the averaged maximum dynamic range of PAClight1_P78A_ (ΔF/F_0_ = 8.28%) vs. PAClight1 (ΔF/F_0_ = 654%) in response to 10μM VIP.

**Extended Data Figure 3.**
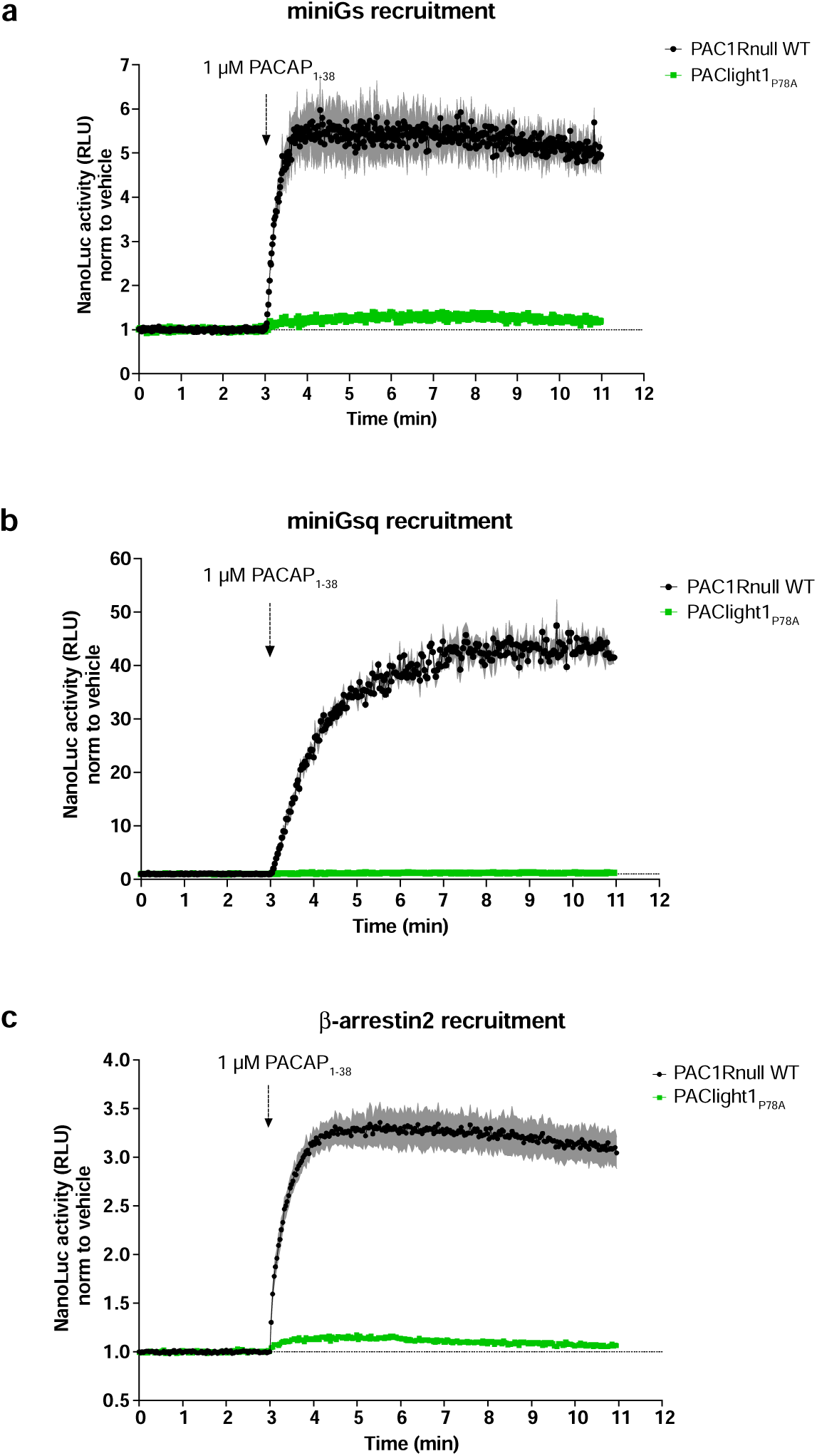
Characterization of PAClight1_P78A_ coupling to transducer proteins. NanoLuc complementation assays were employed to measure the ability of human SmBiT-PAC1R or SmBiT-PAClight1_P78A_ to recruit LgBiT-miniGs (a), -miniGsq (b) or -β-arrestin2 (c) in an agonist-induced manner. Luminescence emission (relative luminescence units, RLU) was monitored during application of agonist (PACAP_1-38_) and was normalized to control cells with application of buffer (vehicle). Dots represent average datapoints, shades represent SEM. n=3 for each condition.

**Extended Data Figure 4.**
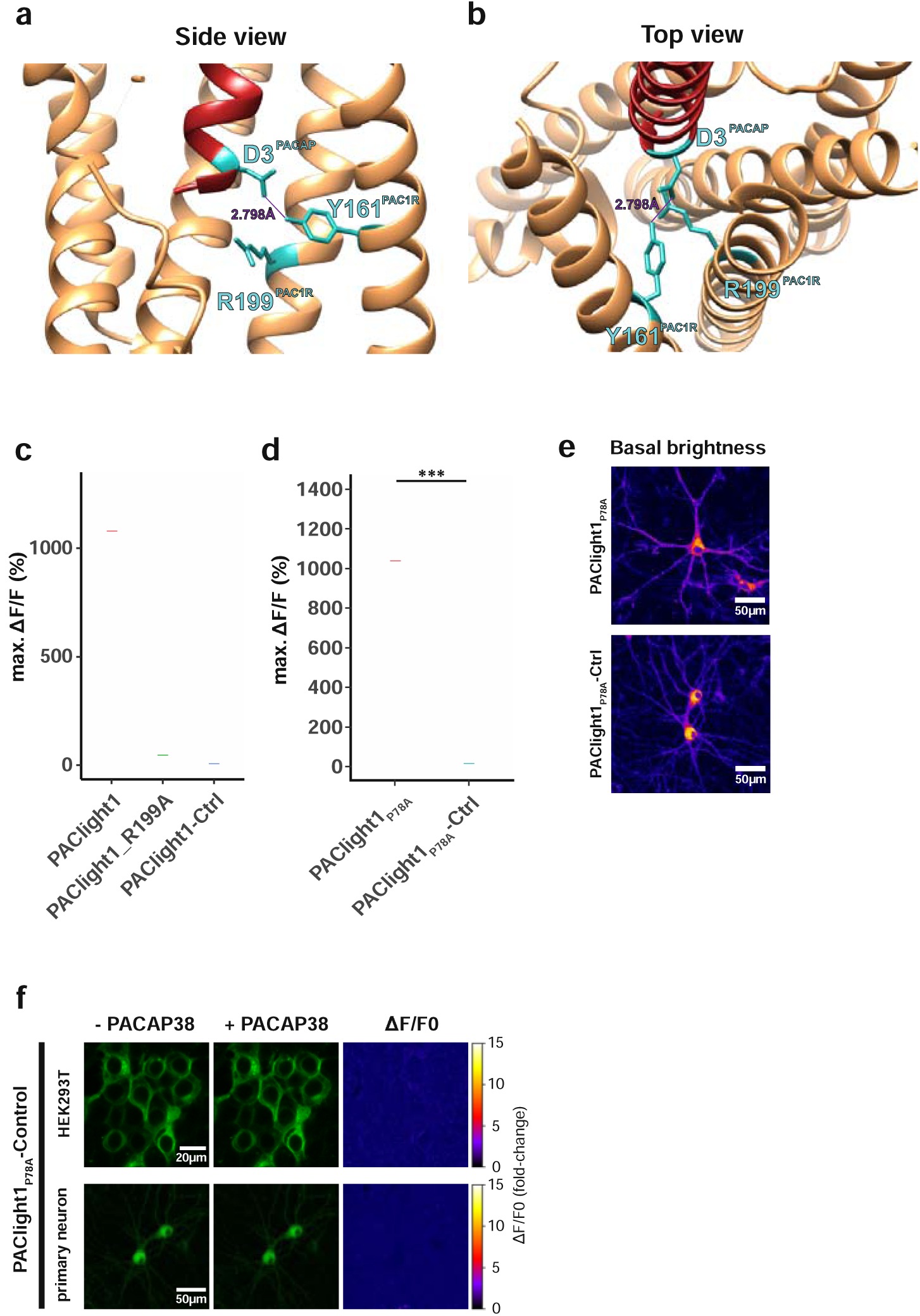
Development of a PAClight1_P78A_-ctrl sensor. **a)** Side view of the protein structure of the hmPAC1R transmembrane domain (beige) in interaction with PACAP_1-38_ (red). Residues Y161 and R199 of the hmPAC1R, as well as residue D3 of PACAP are highlighted in cyan. The hydrogen bond between D3 and Y161 is indicated with a dashed line. **b)** Top view onto the same protein structures depicted in **a)**. **c)** Maximum dynamic range of HEK293T cells expressing PAClight1 (mean= 1075% ΔF/F_0_), PAClight1_R199A (mean= 41.9% ΔF/F_0_), and PAClight1-ctrl (mean= 2.87% ΔF/F_0_). **d)** Maximum dynamic range of HEK293T cells expressing PAClight1_P78A_ (mean= 1034% ΔF/F_0_) and PAClight1_P78A_-ctrl (mean= 10.44% ΔF/F_0_). **e)** Comparison of basal brightness levels of PAClight1_P78A_ and PAClight1_P78A_-ctrl expressed in rat primary neurons via viral transduction. **f)** Representative examples of PAClight1_P78A_-ctrl expressed in HEK293T cells (top row) and rat primary neurons (bottom row) before (left column) and after (middle column) bath application of 10μM PACAP_1-38_. The right column depicts the pixel-wise calculated ΔF/F_0_. No detectable increase in fluorescence is observed.

**Extended Data Figure 5.**
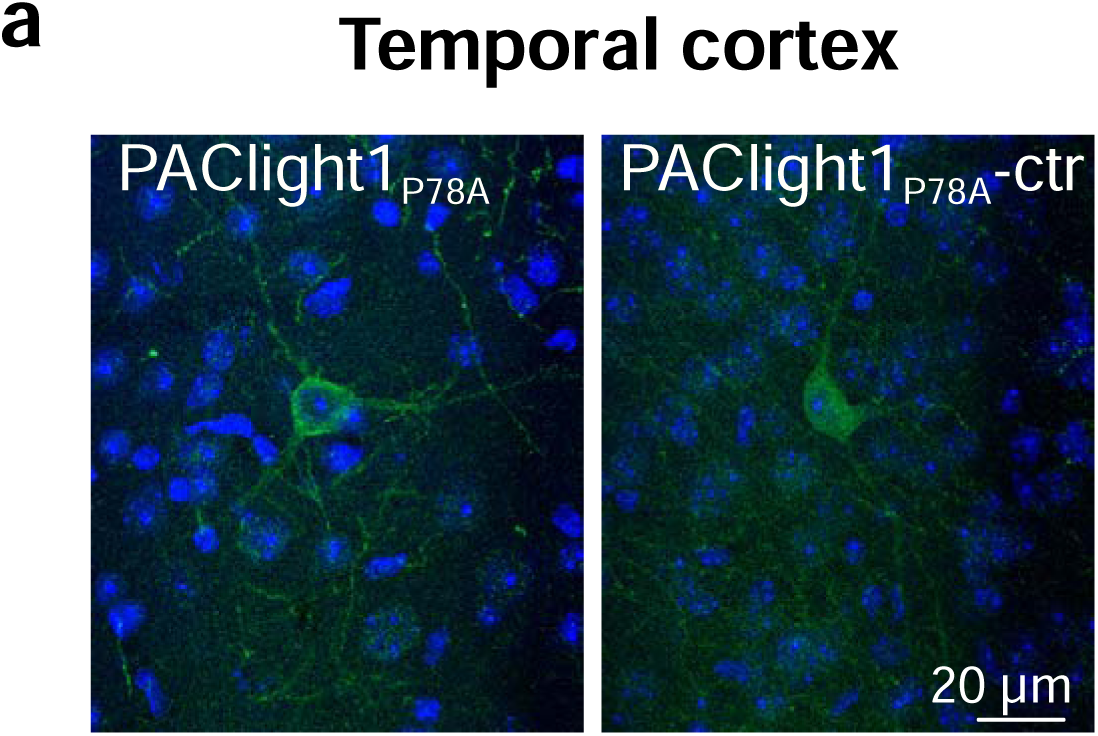
Validation of PAClight1_P78A_ and PAClight1_P78A_-ctrl sensors in mammalian brains. Maximum intensity projections of exemplary confocal images of PAClight1_P78A_ and PAClight1_P78A_-ctrl-expressing neurons in the cortex enhanced with immunostaining and counterstained with DAPI (blue).

